# The TIR-NBS-LRR protein CSA1 is required for autoimmune cell death in Arabidopsis pattern recognition co-receptor *bak1* and *bir3* mutants

**DOI:** 10.1101/2021.04.11.438637

**Authors:** Sarina Schulze, Liping Yu, Alexandra Ehinger, Dagmar Kolb, Svenja C. Saile, Mark Stahl, Mirita Franz-Wachtel, Lei Li, Farid El Kasmi, Volkan Cevik, Birgit Kemmerling

**Affiliations:** ZMBP University of Tübingen, Auf der Morgenstelle 32, 72076 Tübingen, Germany; Interfaculty Institute for Cell Biology, Department of Quantitative Proteomics, University of Tübingen, Auf der Morgenstelle 15, 72076 Tübingen, Germany; Department of Molecular Biology, Max Planck Institute for Developmental Biology, 72076 Tübingen, Germany; The Milner Centre for Evolution, Department of Biology and Biochemistry, University of Bath, Bath BA2 7AY, United Kingdom

**Keywords:** BAK1, cell death, BIR3, CSA1, PTI, ETI, PTI-ETI crosstalk, NLR, receptor kinase, guard

## Abstract

The BRI1-associated kinase BAK1/SERK3 is a positive regulator of multiple leucine rich receptor kinase-mediated signaling pathways including pattern triggered immunity (PTI). Absence or overexpression of BAK1 leads to spontaneous cell death formation. BAK1-interacting receptors (BIR) constitutively interact with BAK1, and plants lacking or overexpressing BIR proteins phenocopy the cell death symptoms observed in *bak1* knock outs or overexpressors. In the interactome of BIR3, the TIR-NBS-LRR protein CONSTITUTIVE SHADE-AVOIDANCE 1 (CSA1) was identified by mass spectrometry. CSA1 physically interacts with BIR proteins and can be detected in complexes with BAK1. Direct interaction was shown only for CSA1 with BIR proteins but not BAK1. Double mutant *bak1 bir3* genotypes develop strong dwarfism and cell death symptoms that are dependent on EDS1 and salicylic acid. Loss of CSA1 blocks *bak1* and *bak1 bir3*-mediated cell death formation thus demonstrating that CSA1 is causal for this type of cell death. We propose that CSA1 guards BIR proteins and initiates autoimmune cell death that is observed when BAK1 BIR complexes are impaired. Our findings reveal how cell death in the absence of BAK1 and BIR3 is executed and links BAK1, a common co-receptor of many pattern recognition receptors, to NLR proteins typically implicated in effector-triggered immunity.

## Introduction

Plants can recognize numerous signals to sense environmental threads and defend themselves against pathogens by perceiving extracellular and intracellular components of pathogens or microbes called microbe-associated molecular patterns (MAMPs) or effectors (Saur et al., 2020; Escocard de Azevedo Manhaes et al., 2021). Extracellular patterns are perceived by cell surface receptors belonging to the receptor kinase (RK) or receptor protein (RP) family (Wan et al., 2019). Effectors that are translocated into the plant cell are perceived by intracellular NOD-like receptors called nucleotide-binding leucine-rich repeat receptors (NLRs). NLRs sense effectors either directly or indirectly by guarding effector targets or effector decoys, mostly proteins important for or involved in immunity (Dangl and Jones, 2001). Both, MAMP and effector recognition leads to defense responses including reactive oxygen species production, salicylate accumulation and transcriptome changes that can fend off pathogens (Lu and Tsuda, 2021). NLRs are divided into the TOLL/INTERLEUKIN RECEPTOR (TIR)-NLRs (TNLs) and coiled-coil domain containing NLRs (CNLs). Downstream components are scarce but TNLs signal via the lipase-like protein ENHANCED DISEASE SUSCEPTIBILITY 1 (EDS1) that functions mutually exclusively either with PHYTOALEXIN DEFICIENT 4 (PAD4) or SENESCENCE ASSOCIATED GENE 101 (SAG101) (Lapin et al., 2020). By contrast, many CNLs signal via the integrin-like protein NON RACE SPECIFIC DISEASE RESISTANCE 1 (NDR1) (Aarts et al., 1998; Knepper et al., 2011). Many effector-sensing NLRs (sensor NLRs) require an additional subclade of NLRs, the RPW8-like helper NLRs that consist of the ACTIVATED DISEASE RESISTANCE 1 (ADR1) and N REQUIREMENT GENES 1 (NRG1) protein families (Jubic et al., 2019). ADR1 proteins can associate with EDS1 and PAD4 while NRG1 proteins only interact with EDS1/SAG101 complexes (Sun et al., 2020).

Leucine-rich repeat receptor kinases (LRR-RK) are the largest subfamily of cell surface receptors and have been assigned functions in many aspects of plant growth, development, and interaction with the environment (Gou et al., 2010). The BRASSINOSTEROID INSENSITIVE 1 (BRI1)-ASSOCIATED KINASE 1 (BAK1) belongs to a five-member LRR-RK subfamily with five LRRs, called SOMATIC EMBRYOGENESIS RECEPTOR KINASES 1 to 5 (SERK1-5) (Hecht et al., 2001). BAK1/SERK3 is a general regulator of other LRR-RKs (Chinchilla et al., 2009) by acting as an interactor and positive regulator of ligand-binding receptors (Chinchilla et al., 2007; Heese et al., 2007; Roux et al., 2011; Ladwig et al., 2015). The best-studied BAK1 interaction partners are FLAGELLIN SENSING 2 (FLS2), which senses bacterial flagellin (or the derived epitope flg22), and BRI1, the major Arabidopsis brassinosteroid (BR) receptor. Biochemical and genetic analyses revealed that BAK1 functions as a co-receptor in both, BRI1 and FLS2 signaling pathways (Li et al., 2002; Nam and Li, 2002; Chinchilla et al., 2007; Heese et al., 2007). Meanwhile, many more RKs and also RPs (via SUPRRESSOR OF BIR1 (SOBIR1)) are linked to the BAK1 co-receptor (Liebrand et al., 2014). The crystal structures of the ligand-bound trimolecular receptor complexes have shown how SERK co-receptors bind to ligand-binding LRR-RKs as well as to the receptor-bound ligands (Santiago et al., 2013; Sun et al., 2013a; Sun et al., 2013b). Ligand-induced association with co-receptors is essential for transmembrane activation of RKs (Song et al., 2016; Hohmann et al., 2017). Subsequent transphosphorylation steps lead to full activation of the cytoplasmic kinase domains and initiation of signaling (Wang et al., 2008; Cao et al., 2013; Bojar et al., 2014).

Reduced levels or overexpression of BAK1 leads to deregulated cell death, indicating that a balanced receptor/co-receptor ratio needs to be maintained to prevent autoimmune cell death (He et al., 2007; Kemmerling et al., 2007; Dominguez-Ferreras et al., 2015). Double mutants of *bak1* and its closest homolog BAK1-LIKE 1 (BKK1)/SERK4 strongly enhance the cell death phenotype of *bak1* single mutants, leading to seedling lethality in double mutant nulls (He et al., 2007). BAK1 also interacts with all members of a family of small LRR-RK called BAK1-INTERACTING RECEPTOR-LIKE KINASES (BIR) which also have effects on cell death control (Gao et al., 2009; Halter et al., 2014; Imkampe et al., 2017; Ma et al., 2017; Hohmann et al., 2018). Four proteins BIR1 to BIR4 form the BIR protein family also named LRR-RK subgroup Xa (Shiu and Bleecker, 2001). Loss-of-function mutants in BIRs have a similar effect on cell death control as described for *bak1* mutants. Furthermore, BIRs are negative regulators of BAK1-mediated immunity. BIRs act by constitutively interacting with BAK1 in the absence of ligands and thus prevent unwanted interaction with ligand-binding receptors. After ligand activation, BIRs are released from the complex and BAK1 can associate with the ligand-bound receptor complex partners. BIR2 affects flg22- and elf18-induced signaling, as well as cell death control, but not BR signaling, while BIR3 has additionally an effect on BL signaling (Halter et al., 2014; Imkampe et al., 2017).

Modelling of the receptor complex revealed that a tripartite complex with BAK1, BRI1 and BIR3 is unlikely as BIR3 and BRI1 have a strong preference for the same binding site on BAK1 (Grosseholz et al., 2020). This is supported by the crystal structure of the extracellular domains of the BIR1 BAK1 complex that revealed an interaction interface that is very similar to the one that FLS2 uses for binding to BAK1 and therefore supports a competition of ligand binding receptors and BIR proteins for BAK1 (Ma et al., 2017; Hohmann et al., 2018).

Double mutant *bak1 bir3* plants show a severe dwarf phenotype that is associated with spontaneous cell death, mimicking the phenotype of *bak1 bkk1* mutants described above (Imkampe et al., 2017). Information on how *bak1* and *bir-*mediated cell death is initiated and which pathway components are involved is still limited. A number of suppressor screens have been performed and only a few suppressors of *bak1 bkk1* or *bir1*-mediated cell death have been published so far. These include glycosylation factors, ER quality control components, and nuclear import and export proteins (de Oliveira et al., 2016; Du et al., 2016). The *bir1*-mediated cell death is suppressed by *sobir1, bak1* and *pad4* mutation (Gao et al., 2009; Liu et al., 2016). Controversial outcomes have been reported for the rescue of *bak1 bkk1*-mediated cell death by *pad4* or *ndr1* mutation (de Oliveira et al., 2016; Gao et al., 2017). The *bak1 bkk1* mutants cell death is rescued also by loss-of-function mutants of the cyclic nucleotide gated ion channel 20 (CNGC20) (Yu et al., 2019) and the nucleoporin *suppressor of bak1 bkk1* (SBB1) mutant, a protein that controls nucleocytoplasmic trafficking (Du et al., 2016). MODIFIER OF SNC1 3 (MOS3) is a member of the same nuclear envelope complex and was identified as a nucleoporin that affects multiple disease resistance pathways. *MOS3* mutation also suppresses *bak1 bkk1-*mediated cell death (Zhang and Li, 2005).. Recently Wu et al. (2020) revealed the helper NLR ADR1 family as part of the *bak1 bkk1* cell death pathway. These data indicate that NLR signaling is involved in *bak1*- and maybe also *bir*-mediated cell death. But none of the screens have identified the guard or the specific signal that is needed for the cell death phenotypes.

Further evidence for a specific involvement of NLR-type resistance proteins in *bak1 bir3-*mediated cell death comes from the mass spectrometry based interactome search of BIR3 where we identified the TNL CONSTITUTIVE SHADE AVOIDANCE 1 (CSA1) as a potential interactor of BIR3 (Faigon-Soverna et al., 2006). Our data show that BIR3 can interact with CSA1 and that *bak1* and *bak1 bir3-*mediated cell death is blocked by mutation in *csa1.* This suggests that lack of wildtype (wt) BAK1 and/or BIR proteins activate CSA1 leading to cell death, indicating that functional integrity of the BAK1 BIR protein complex is guarded by CSA1.

## Results

### Cell death in *bak1 bir3* double mutant is dependent on EDS1, SA and NRGs

To investigate how the absence of BAK1 and BIR3 initiates cell death we checked the potential immunity-related cell death pathways known in plants and created triple and quadruple mutants with components of these pathways. We crossed plants expressing a salicylic acid degrading bacterial enzyme NahG and mutants of known ETI pathway components such as *eds1, pad4* and *sag101* with *bak1 bir3* mutants. The *eds1* mutant can partially block *bak1 bir3*-mediated growth phenotypes (Figure 1A), and after *Alternaria brassicicola* infection cell death is reduced to wt levels and infection symptoms are as low as in the hyperresistant *eds1* mutant (Figure 1B-D). The effect of *eds1* mutation on the *bak1 bir3* phenotype suggests that a TNL might be involved in guarding the BAK1 BIR3 complex and initiate cell death when one or both of the components are absent. *pad4 and sag101* mutants had only a weak effect on the *bak1 bir3* growth phenotype (Supplemental Figure1). This indicates that the EDS1 SAG101 or EDS1 PAD4 hubs in NLR-mediated immunity downstream of TNLs are redundantly and at least partially necessary for *bak1 bir3*-mediated cell death. SA is indispensable for *bak1*-mediated cell death (Gao et al., 2017). Expression of *NahG,* that degrades salicylic acid, also affects *bak1 bir3*-mediated cell death (Supplemental Figure1). Our findings show that wt BAK1 and BIR3 contain cell death that is executed by EDS1-dependent complexes and SA, leading to autoimmunity-associated runaway cell death in plants lacking a functional BAK1 BIR3 complex. This suggests that components of the TNL-mediated ETI pathway contribute to and are necessary for *bak1 bir3*-mediated cell death, but the requirement of additional components for the observed phenotypes also needs to be postulated.

**Figure 1:**
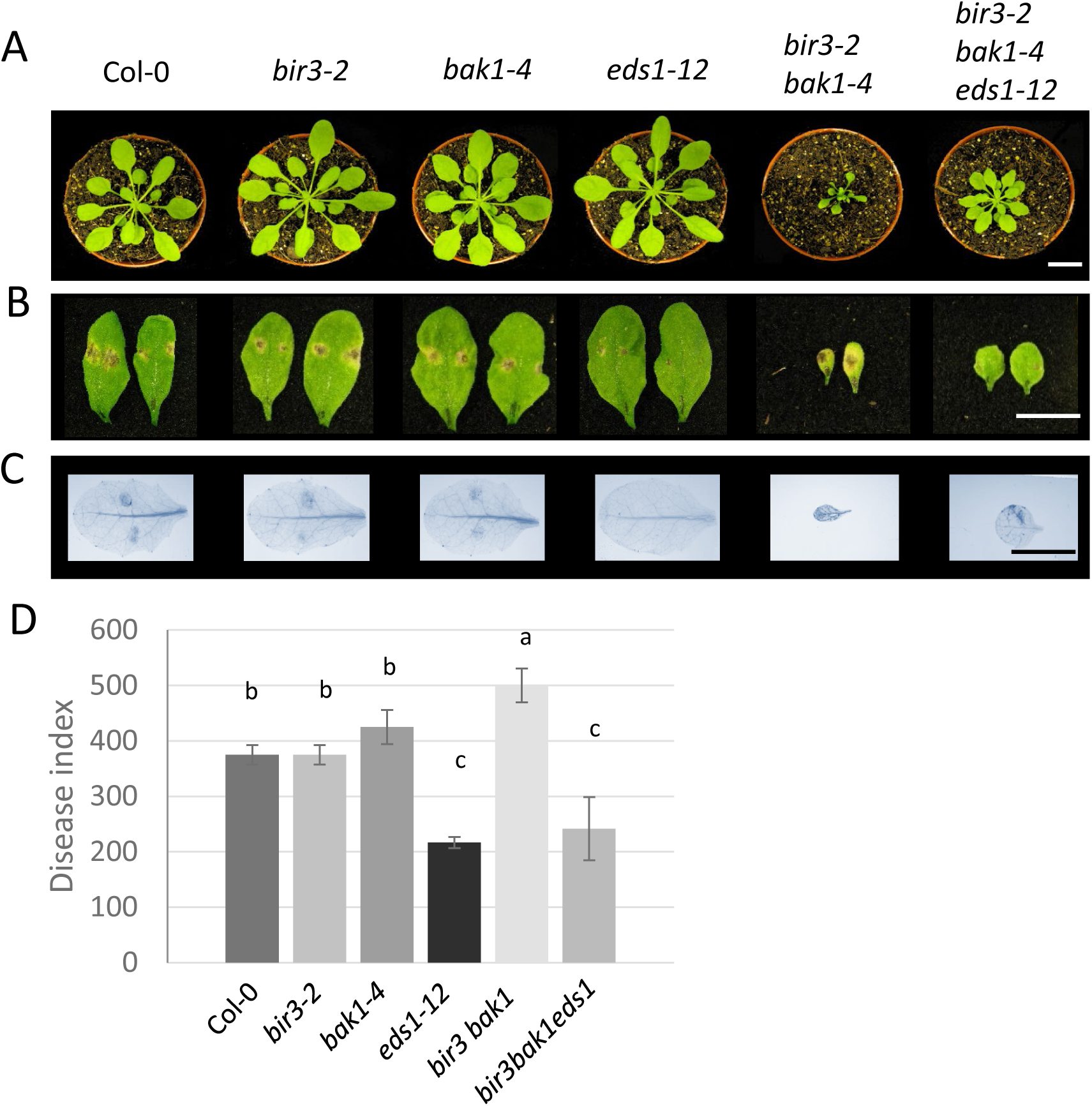
Loss-of *EDS1* can partially block *bak1 bir3* induced cell death. **(A)** Representative pictures of the morphological phenotypes of 6-week-old Col-0, *bir3-2, bak1-4, eds1-12*, the double mutant *bak1-4 bir3-2* and the triple mutant *bak1-4 bir3-2 eds1-12* are shown. **(B)** *Alternaria brassicicola* droplet-infected leaves of the genotypes shown in (A) 13 days after inoculation. **(C)** Leaves of the same genotypes as in (A) and (B) droplet-infected with *Alternaria brassicicola* and trypan blue stained. The scale bar in (A), (B) and (C) represents 10 mm. **(D)** Disease indices of *Alternaria brassicicola* infected leaves of the indicated genotypes 13 days after infection shown as mean ± SE (n=12). Different letters indicate significant differences according to one-way ANOVA and Tukey’s HSD test (p<0.05). The experiments were repeated at least three times with similar results.

Lapin et al. (2019) postulated that EDS1-SAG101 and NRG1s co-evolved as a functional TNL-dependent cell-death and resistance module. Since we observed a partial suppression of the *bak1 bir3* phenotype by *eds1*, we determined whether the NRG1 family is also involved in mediating *bak1 bir3* cell death. Therefore, we crossed *nrg1.1 nrg1.2* double mutants with *bak1 bir3* double mutants. The *nrg1.1 nrg1.2 bak1 bir3* mutants are larger than *bak1 bir3* mutants showing that NRG1s, as part of the TNL cell death pathway, are also contributing to the *bak1 bir3* phenotype (Supplemental Figure 2).

### The interactome of BIR3 contains the TNL CSA1

We determined the *in vivo* interactome of BIR3-YFP by liquid chromatography-electron spray ionization tandem mass spectrometry (LC-ESI-MS/MS) in Arabidopsis plants. MS analyses revealed BAK1 and other SERKs as the most abundant interaction partners of BIR3 (Supplemental Table 1), confirming the strong interaction with BAK1 published previously (Xing et al., 2007; Gao et al., 2009; Halter et al., 2014; Imkampe et al., 2017). In addition, other known RKs such as SOBIR1, FERONIA and MIK2 have been detected, indicating that BIR3 interacts with multiple known but also yet undescribed RKs (Liebrand et al., 2014; Stegmann et al., 2017; Van der Does et al., 2017). Thus, BIR3 seems to be a general interactor of RKs and therefore likely also affects multiple signaling pathways (Supplemental Table 1).

From the BIR3-interacting protein candidates we identified a protein with a unique peptide sequence LPDSLGQLK of a known NLR as a potential candidate for executing cell death in *bak1 bir3* genotypes. (Supplemental Figure 3A). This peptide corresponds to the TNL protein CONSTITUTIVE SHADE AVOIDANCE 1 (CSA1) that was previously shown to affect shade avoidance (Faigon-Soverna et al., 2006) and autoimmunity (Xu et al., 2015) (Supplemental Figure 3 B, C).

CSA1 is indispensable for the autoimmune phenotypes initiated by hyperactive *CHILLING SENSITIVE 3 (CHS3)* alleles, an NLR-encoding gene localized adjacent to *CSA1* on chromosome 5. CSA1 and CHS3 are proposed to act together as a sensor and executer pair of NLRs (Xu et al., 2015; Castel et al., 2019; Wu et al., 2019). The identification of CSA1 in our BIR3-interactome and the partial suppression of the *bak1 bir3* phenotype by components of TNL-mediated immunity strongly suggest a potential involvement of CSA1 in the *bak1 bir3* cell death phenotype.

### BIR3 interacts with CSA1

To confirm the interaction of CSA1 and BIR3 identified *in planta* by MS analysis, co-immunoprecipitation (co-IP) experiments were performed. BIR3-GFP and CSA1-V5 fusion proteins were expressed under the constitutively active 35S promoter in Nicotiana (*N.) benthamiana*. For the expression of respective proteins, p19, a silencing suppressor, was co-infiltrated in order to suppress the RNA silencing machinery of the plant. Leaves infiltrated with p19 only served as negative controls to detect unspecific binding of proteins to the beads used for immunoprecipitation. Expression of CSA1-V5 and BIR3-GFP alone served as additional negative controls, to ensure that no unspecific binding to the beads is detected instead of proteins co-immunoprecipitated with BIR3-GFP. CSA1 is detectable in the co-immunoprecipitated samples but not in control samples, suggesting that CSA1 can interact with BIR3 (Figure 2). Interaction studies in Arabidopsis turned out to be limited due to technical and biological difficulties. Expression of genomic constructs of CSA1-V5 result in low expressing or lethal plants, the BIR3 antibody has a low affinity and is insufficient to detect co-immunoprecipitated BIR3 and the amount of expressed and detectable immunoprecipitated CSA1 was too little to prove interaction. The other expressed BIR family members BIR1 and 2 can also interact with CSA1 in co-IP after transient expression in *N. benthamiana* (Supplemental Figure 4).

**Figure 2:**
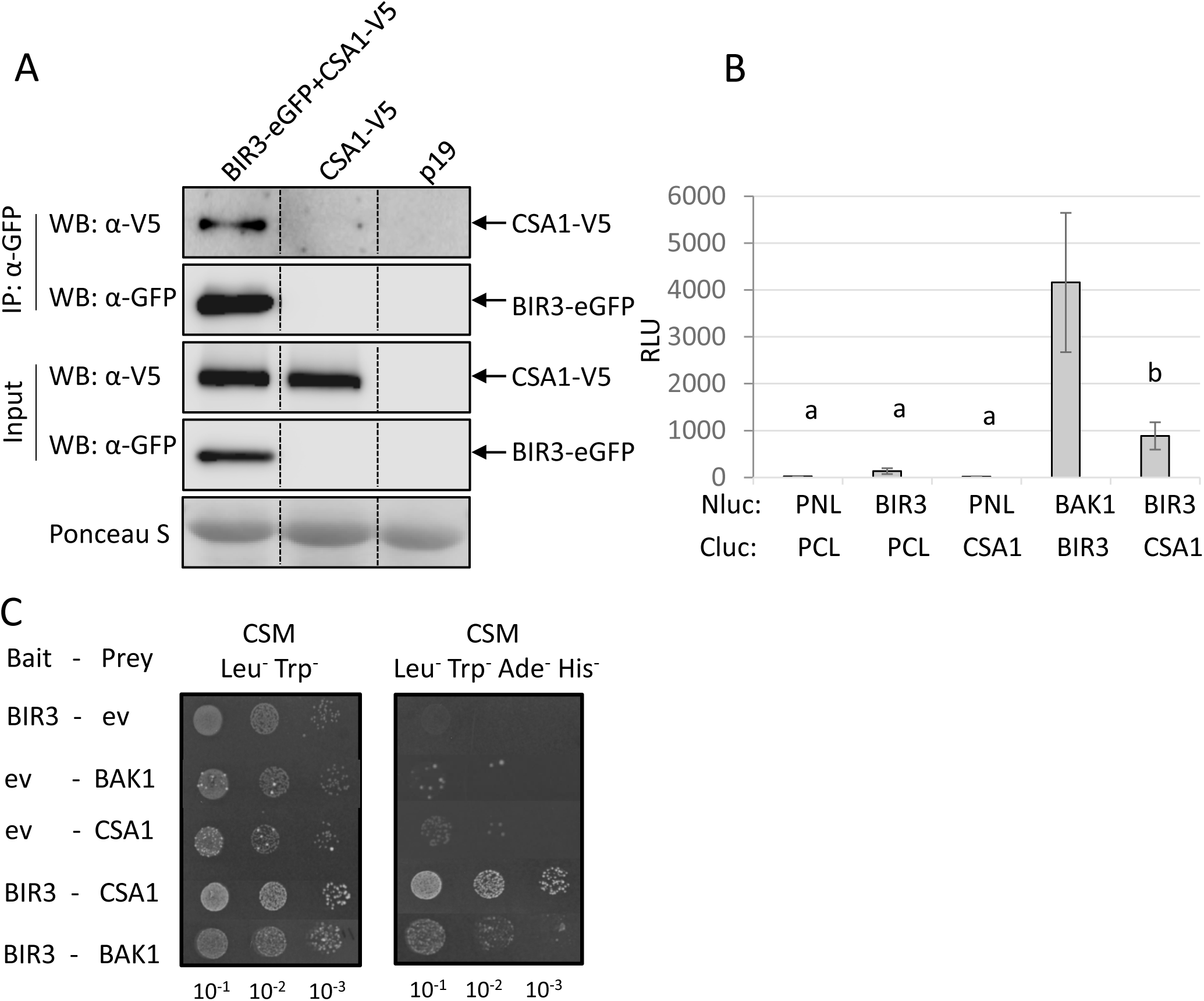
CSA1 can interact with BIR3. **(A)** Western Blots after co-immunoprecipitation with GFP-traps of transiently in *N. benthamiana* expressed BIR3-eGFP and CSA1-V5 detected with α-GFP and α-V5 antibodies are shown. Protein input is shown by Western blot analyses of protein extracts before IP with antibodies against the respective tags. p19 is a silencing inhibitor expressed alone as a background control. Ponceau S staining shows protein loading. Dotted lines indicate cut and rearranged parts of the same blot. **(B)** Split-Luciferase assays with transiently expressed BIR3-Nluc and CSA1-Cluc fusion proteins show reconstituted luciferase activity measured in relative light units (RLU) indicating that the two proteins are in close vicinity. BAK1-Nluc and BIR3-Cluc constructs serve as positive controls. Empty vector controls (PNL, PCL) serve as negative controls. **(C)** Split-ubiquitin yeast growth assays containing the two indicated proteins fused to N- and C-terminal parts of ubiquitin were performed (ev, empty vector). Yeast was grown at three different 1 to 10 dilutions on medium selecting for vector transformation (CSM -Leu^−^, Trp^−^) and for interaction (CSM-Leu^−^, Trp^−^, Ade^−^, His^−^). Growth was monitored after 1 d for the vector-selective control plates and after 3 d for the interaction plates. BIR3 and BAK1 serve as positive controls, empty vector controls as negative controls. All experiments were repeated at least three times with similar results.

Localization studies with fractionated plant extracts and confocal laser scanning microscopy revealed that CSA1 is predominantly localized to microsomal fraction and the plasma membrane, respectively (Supplemental Figure 5), which is in agreement with interaction of plasma membrane-resident BIR3.

To support these data, BIR3 and CSA1 coding sequences fused to the N- and C-terminal parts of Luciferase (Luc) were transiently expressed *in planta* (*N. benthamiana*) and tested for restoration of luciferase enzyme activity. The increase in luciferase activity, as quantified by relative light units emitted from degraded luciferin, shows that the luciferase can reconstitute when CSA1-NLuc and BIR3-CLuc are expressed together but not when expressed with the empty vector controls, confirming that also in this experimental setup CSA1 and BIR3 are interacting *in planta* (Figure 2 B; Supplemental Figure 6).

### The interaction of CSA1 and BIR3 is direct

While proteins detected in co-IPs do not necessarily interact directly, the Split-LUC assays already suggests a very close vicinity of the proteins. If this is indeed a direct interaction was tested with the Split-ubiquitin assay (SUS) in yeast. CSA1 and BIR3 were fused to N- and C-terminal parts of ubiquitin and expressed in yeast. The growth rescue on limiting medium strongly suggests that CSA1 can directly interact with BIR3 but not with BAK1 (Figure 2 C; Supplemental Figure 7). We hypothesize that CSA1 directly guards BIR3 and that cell death in *bak1* mutants may be mediated via BIR3 by CSA1.

### The CSA1 partner CHS3 does not interact directly with BIR3

CSA1 works as a pair with its genetic neighbor the TNL protein CHILLING SENSITIVE 3 (CHS3) (Xu et al., 2015; Van de Weyer et al., 2019). In a suppressor screen for auto-active *chs3-2D* mutants CSA1 was found to be indispensable for the execution of the autoimmune cell death exhibited by the auto-active *chs3-2D* allele (Xu et al., 2015). CHS3 is a TNL with additional integrated domains (LIM (<u>L</u>in-11, Isl-1 and <u>M</u>ec-3 domain) domain and putative DA-1 protease domain) at the C-terminus (Yang et al., 2010) and encoded in a head-to-head orientation with *CSA1* (Supplemental Figure 8). Furthermore, CSA1 and CHS3 were found to specifically associate in Arabidopsis as well as in *N. benthamiana* plants (Parkes, 2020). We detected CHS3 in co-IPs with BIR3 when both proteins were transiently expressed in *N. benthamiana* (Figure 3 A). BIR1 and BIR2 can also interact in co-IPs with CHS3 (Supplemental Figure 9). In all assays the interaction appeared weaker than the CSA1 BIR3 interaction. The BIR3 CHS3 interaction was neither confirmed in Split-Luc assays nor in SUS assays indicating that CSA1 is the direct interactor and that CHS3 might be part of the same complex but not in physical contact with BIR3 (Figure3 B, C; Supplemental Figure 6, 7).

**Figure 3:**
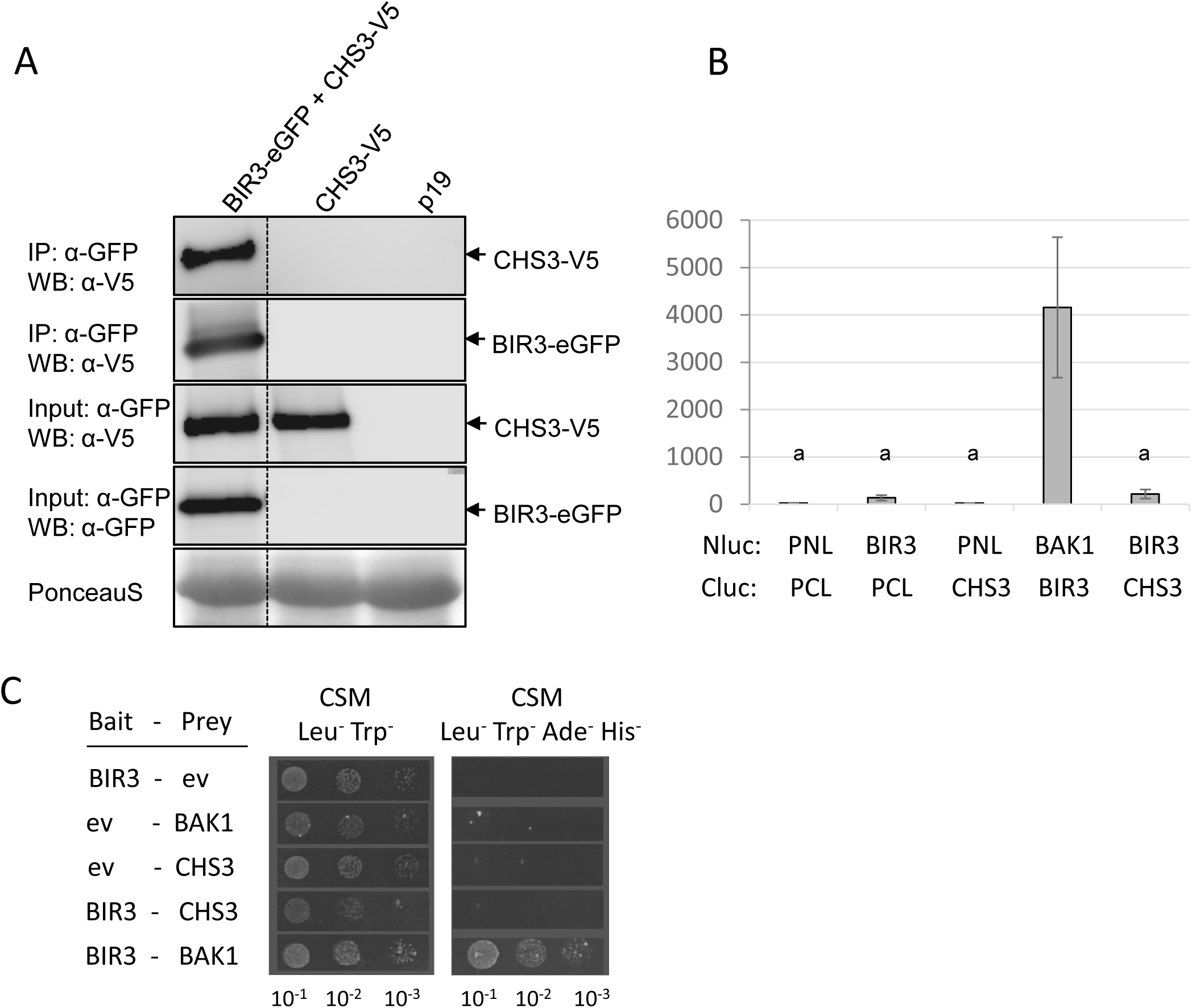
The CSA1 partner CHS3 resides in the BIR3 complex but does not directly interact with BIR3. **(A)** Western Blots after co-immunoprecipitation with GFP-traps of transiently in *N. benthamiana* expressed BIR3-eGFP and CHS3-V5 protein detected with α-GFP and α-V5 antibodies are shown. Protein input is shown by Western blot analysis of protein extracts before IP with antibodies against the respective tags. p19 is a silencing inhibitor expressed alone as a background control. Ponceau S staining shows protein loading. Dotted lines indicate cut and rearranged parts of the same blot. **(B)** Split-Luc assays with transiently expressed BIR3-Nluc and CHS3-Cluc fusion proteins were analyzed for reconstituted luciferase activity measured in relative light units (RLU). BAK1-Nluc and BIR3-Cluc constructs serve as positive controls. Empty vector controls (PNL, PCL) serve as negative controls. Different letters indicate significant differences according to one-way ANOVA and Tukey’s HSD test (p<0.05). **(C)** Split-ubiquitin yeast growth assays containing the two indicated proteins fused to N- and C-terminal parts of ubiquitin were performed (ev, empty vector). Yeast was grown at three different 1 to 10 dilutions on medium selecting for vector transformation (CSM -Leu^−^, Trp^−^) and for interaction (CSM-Leu^−^, Trp^−^, Ade^−^, His^−^). Growth was monitored after 1 d for the vector-selective control plates and after 3 d for the interaction plates. BIR3 and BAK1 serve as positive controls, empty vector controls as negative controls. All experiments were repeated at least three times with similar results.

### CSA1 and CHS3 are in complex with BAK1

If CSA1 and CHS3 are in complexes with BIR3 the question arises if they also interact with BAK1. BAK1-GFP and CSA1 can be detected in the same co-immunoprecipitated complexes, when transiently expressed in *N. benthamiana* (Figure 4 A). However, we failed to confirm this interaction in Split-luciferase and SUS assays (Fig 4 B, C). These results indicate that BAK1 and CSA1 might be part of the same complex, but they most likely do not physically interact. Weak interaction of CHS3 with BAK1 was also detectable in co-IPs of transiently expressed tagged proteins, but neither in Split-Luc assays nor in SUS assays a close vicinity or direct interaction could be shown (Figure 5), similar to the BAK1 CSA1 association. Subcellular localization assays show that CHS3 is localized to the plasma membrane but also to the soluble fraction and can be localized in the nucleus, which is in agreement with a weaker interaction with membrane-resident RKs (Supplemental Figure 5).

**Figure 4:**
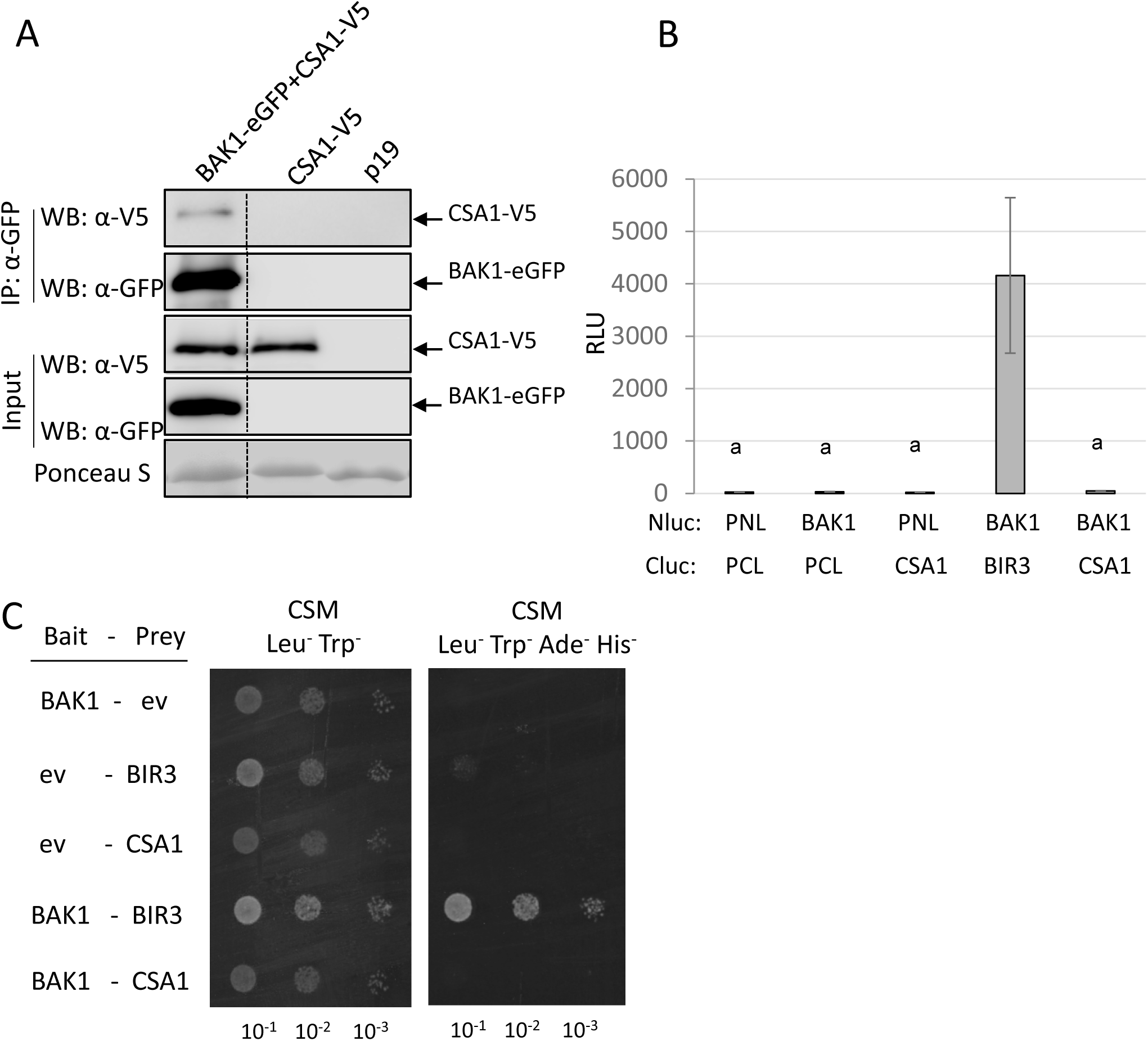
CSA1 is in the same complex with BAK1 but the interaction is not direct. **(A)** Western Blots after co-immunoprecipitation with GFP-traps of transiently in *N. benthamiana* expressed BAK1-eGFP and CSA1-V5 detected with α-GFP and α-V5 antibodies are shown. Protein input is shown by Western blot analysis of protein extracts before IP with antibodies against the respective tags. p19 is a silencing inhibitor expressed alone as a background control. Ponceau S staining shows protein loading. Dotted lines indicate cut and rearranged parts of the same blot. **(B)** Split-Luc assays with transiently expressed BAK1-Nluc and CSA1-Cluc fusion proteins were analyzed for reconstituted luciferase activity measured in relative light units (RLU). BAK1-Nluc and BIR3-Cluc constructs serve as positive controls. Empty vector controls (PNL, PCL) serve as negative controls. Different letters indicate significant differences according to one-way ANOVA and Tukey’s HSD test (p<0.05). **(C)** Split-ubiquitin yeast growth assays containing the two indicated proteins fused to N- and C-terminal parts of ubiquitin were performed (ev, empty vector). Yeast was grown at three different 1 to 10 dilutions on medium selecting for vector transformation (CSM -Leu^−^, Trp^−^) and for interaction (CSM-Leu^−^, Trp^−^, Ade^−^, His^−^). Growth was monitored after 1 d for the vector-selective control plates and after 3 d for the interaction plates. BIR3 and BAK1 serve as positive controls, empty vector controls as negative controls. All experiments were repeated at least three times with similar results.

**Figure 5:**
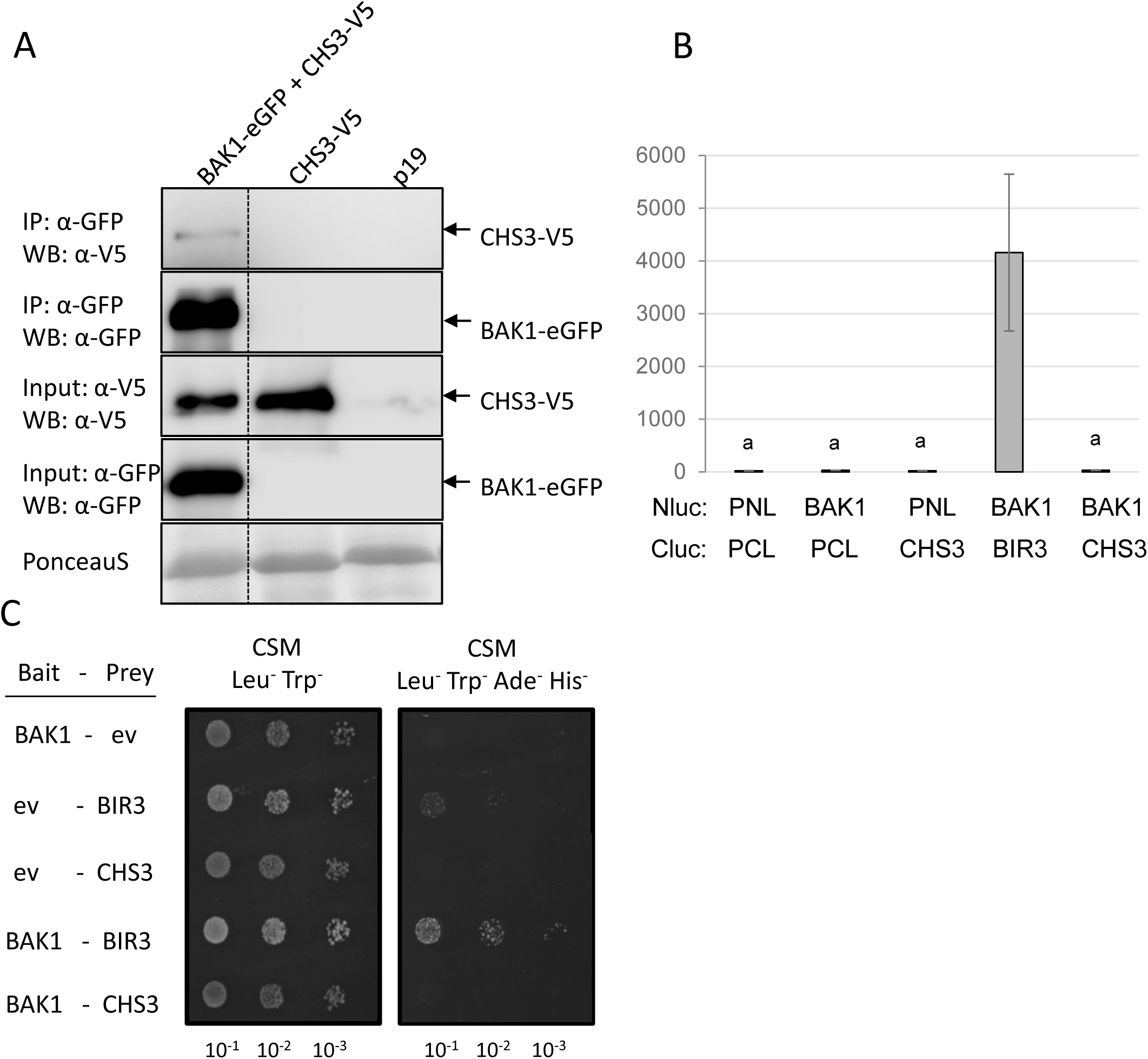
CHS3 is in the same complex with BAK1 but the interaction is not direct. **(A)** Western Blots after co-immunoprecipitation with GFP-traps of transiently in *N. benthamiana* expressed BAK1-eGFP and CHS3-V5 detected with α-GFP and α-V5 antibodies are shown. Protein input is shown by Western blot analysis of protein extracts before IP with antibodies against the respective tags. p19 is a silencing inhibitor expressed alone as a background control. Ponceau S staining shows protein loading. Dotted lines indicate cut and rearranged parts of the same blot. **(B)** Split-Luc assays with transiently expressed BAK1-Nluc and CHS3-Cluc fusion proteins were analyzed for reconstituted luciferase activity measured in relative light units (RLU). BAK1-Nluc and BIR3-Cluc constructs serve as positive controls. Empty vector controls (PNL, PCL) serve as negative controls. Different letters indicate significant differences according to one-way ANOVA and Tukey’s HSD test (p<0.05). **(C)** Split-ubiquitin yeast growth assays containing the two indicated proteins fused to N- and C-terminal parts of ubiquitin were performed (ev, empty vector). Yeast was grown at three different 1 to 10 dilutions on medium selecting for vector transformation (CSM -Leu^−^, Trp^−^) and for interaction (CSM-Leu^−^, Trp^−^, Ade^−^, His^−^). Growth was monitored after 1 d for the vector-selective control plates and after 3 d for the interaction plates. BIR3 and BAK1 serve as positive controls, empty vector controls as negative controls. All experiments were repeated at least three times with similar results.

Taken together, these data indicate that CSA1 guards BIR proteins by directly interacting with them and CHS3 and BAK1 are partners in CSA1 complexes but not direct interaction partners in this complex. Whether BAK1 and CHS3 are simultaneously in complex with CSA1 needs to be studied in future experiments.

### CSA1 is required for *bak1 bir3-*mediated cell death

Loss of *BAK1* and *BIR3* causes strong autoimmune cell death in Arabidopsis (Imkampe et al., 2017). As we identified CSA1 in the interactome of BIR3, we generated a *bak1 bir3 csa1* triple mutant to test whether loss of CSA1 can block *bak1 bir3*-initiated cell death. The triple mutants are significantly larger, fertile and less affected in autoimmune cell death (Figure 6 A). The cell death and spreading necrosis observed in *bak1* and *bak1 bir3* mutants upon *Alternaria (A.) brassicicola* infections is inhibited in *bak1 bir3 csa1* triple mutants (Figure 6 C,D). SA levels that are strongly enhanced in *bak1 bir3* double mutants are reverted in the *bak1 bir3 csa1* triple mutant to levels detectable in the single mutant parents (Figure 6 E). PAMP-induced ROS burst is not affected in *csa1* mutants while basal and PAMP-induced *FRK1* gene expression that is strongly induced in *bak1 bir3* double mutants is lowered to single *bak1* mutant levels in the *bak1 bir3 csa1* triple mutant (Figure 6 F,G) indicating that CSA1-mediated cell death responses potentiate flg22-triggered immune responses. The *bir1* dwarfism phenotype is not affected by *csa1* mutation indicating that *bir1* and *bak1 bir3* cell death are not identical and require different downstream components (Supplemental Figure 10).

**Figure 6:**
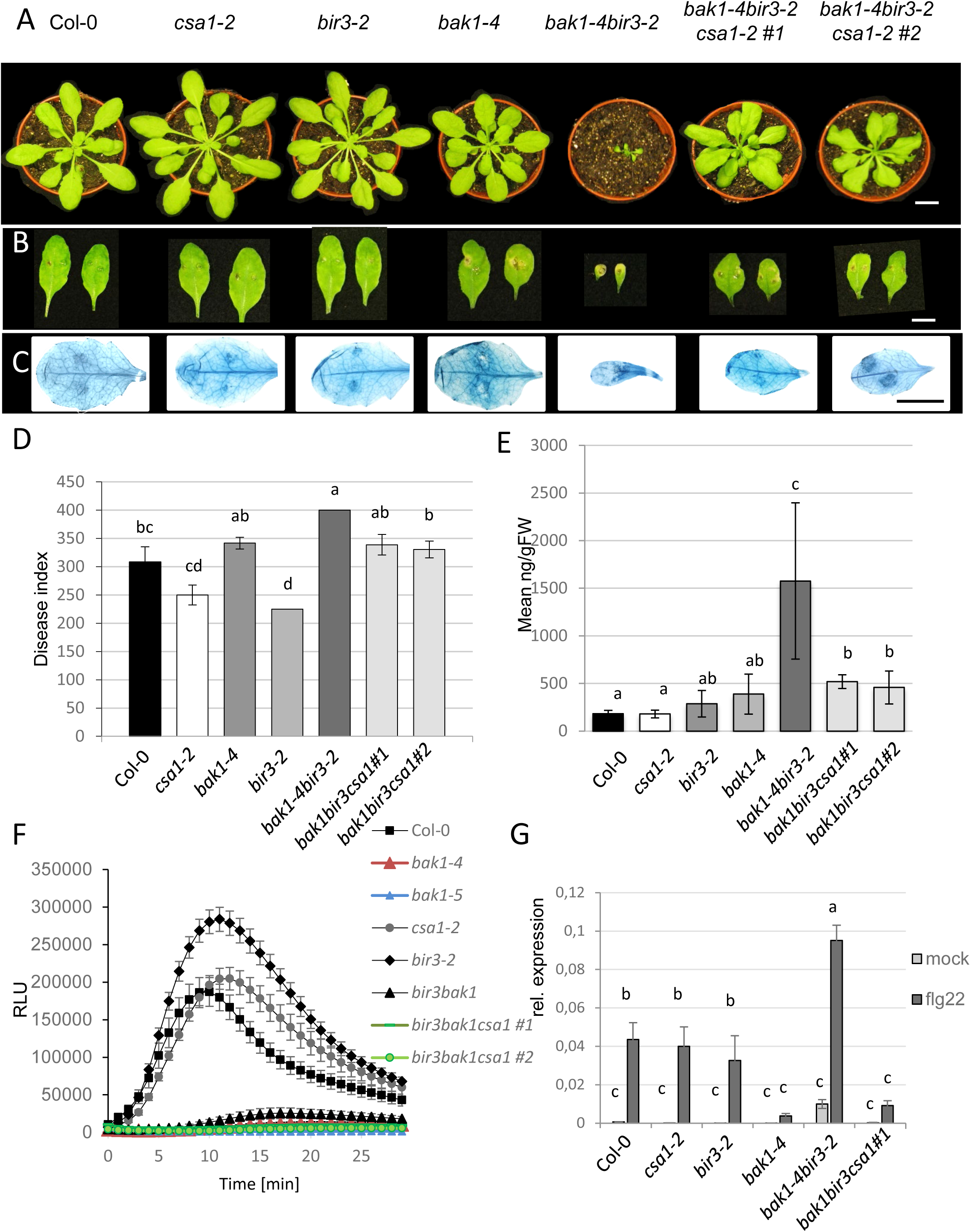
CSA1 is necessary for *bak1 bir3*-mediated cell death. **A)** Representative pictures of the morphological phenotypes of 6-week-old plants of the indicated genotypes are shown. **(B)** *Alternaria brassicicola* infected leaves of the genotypes shown in (A) 13 days after inoculation. **(C)** Infected leaves of the genotypes shown in (A, B) stained with trypan blue for cell death. The scale bar in (A), (B) and (C) represents 10 mm. **(D)** Disease indices of *Alternaria brassicicola* infected leaves of the indicated genotypes 13 days after infection shown as mean ± SE (n=12). **(E)** Total salicylic acid levels measured in untreated leaf material of plants of the indicated genotypes relative to the fresh weight. **(F)** ROS production was measured as relative light units (RLU) in a luminol based assay. Leaf pieces of plants of the indicated genotypes were elicited with 100 nM flg22 and ROS production was measured over a period of 30 min. Values are mean ± SE (n=9). **(G)** *FRK1* marker gene expression in leaves of plants of the indicated genotypes was measured by qRT-PCR analysis 3 hours after 100 nM flg22 or mock treatment. *FRK1* expression was normalized to *EF1*α. Results are mean ± SE (n=9). Different letters indicate significant differences according to one-way ANOVA and Tukey’s HSD test (p<0.05). The experiments were repeated at least three times with similar results.

Crosses of *bak1 bir3* with *chs3* mutants revealed that CHS3 is less involved in the growth phenotype as is CSA1 (Supplemental Figure 11 A). However, loss of CHS3 in the *bak1 bir3* background also affects the autoimmune cell death shown by trypan blue staining of uninfected plants and the cell death induced by *Alternaria* infections (Supplemental Figure 11 B,C). This suggests that CHS3 is partially contributes to the autoimmune cell death symptoms and the development of spreading cell death after *Alternaria* infections in the *bak1 bir3* mutant (Supplemental Figure 11 C, D).

Altogether, these data show that CSA1 and, to a lesser extent, also CHS3 are necessary for triggering autoimmune cell death in *bak1 bir3* mutants. The TNL CSA1 interacts with BIR3 and induces cell death reactions in the absence of BAK1 and/or BIR3 and thus acts as a guard of functional, active BIR3.

### CSA1 is required for *bak1*-mediated cell death

If CSA1 can only interact with BIR3 directly and not with BAK1 does it also guard BAK1 and mediates *bak1* loss of function cell death? We crossed *csa1* also to *bak1* alone and monitored the macroscopic phenotype and cell death. Macroscopically *bak1 csa1* mutants are similar to *bak1* single mutants (Figure 7A). As the *bak1* mutants show only weak spontaneous cell death without any trigger, we infected the indicated genotypes with *A. brassicicola* and scored the disease index. In the double mutant *bak1 csa1*, both, disease indices and visible symptoms are significantly weaker as compared to those in *bak1* single mutants and resemble the phenotype of *csa1* mutants that are slightly less affected than the Col-0 wt plants (Figure 7 B-E). PAMP responses, for example ROS burst after flg22 treatment and *FRK1* gene expression, of the *bak1 csa1* mutants are identical to those observed in *bak1* single mutant responses, suggesting that CSA1 is necessary for the cell death responses triggered in *bak1* mutants but not for the PAMP responses (Figure 7 F, G). Complementation of the *bak1 csa1* mutants and the *bak1 bir3 csa1* mutants with a genomic construct expressing CSA1 under its own promoter complement mutant phenotypes and restore strong cell death symptoms typically observed in *bak1* and *bak1 bir3* mutants (Supplemental Figure 12, 13).

**Figure 7:**
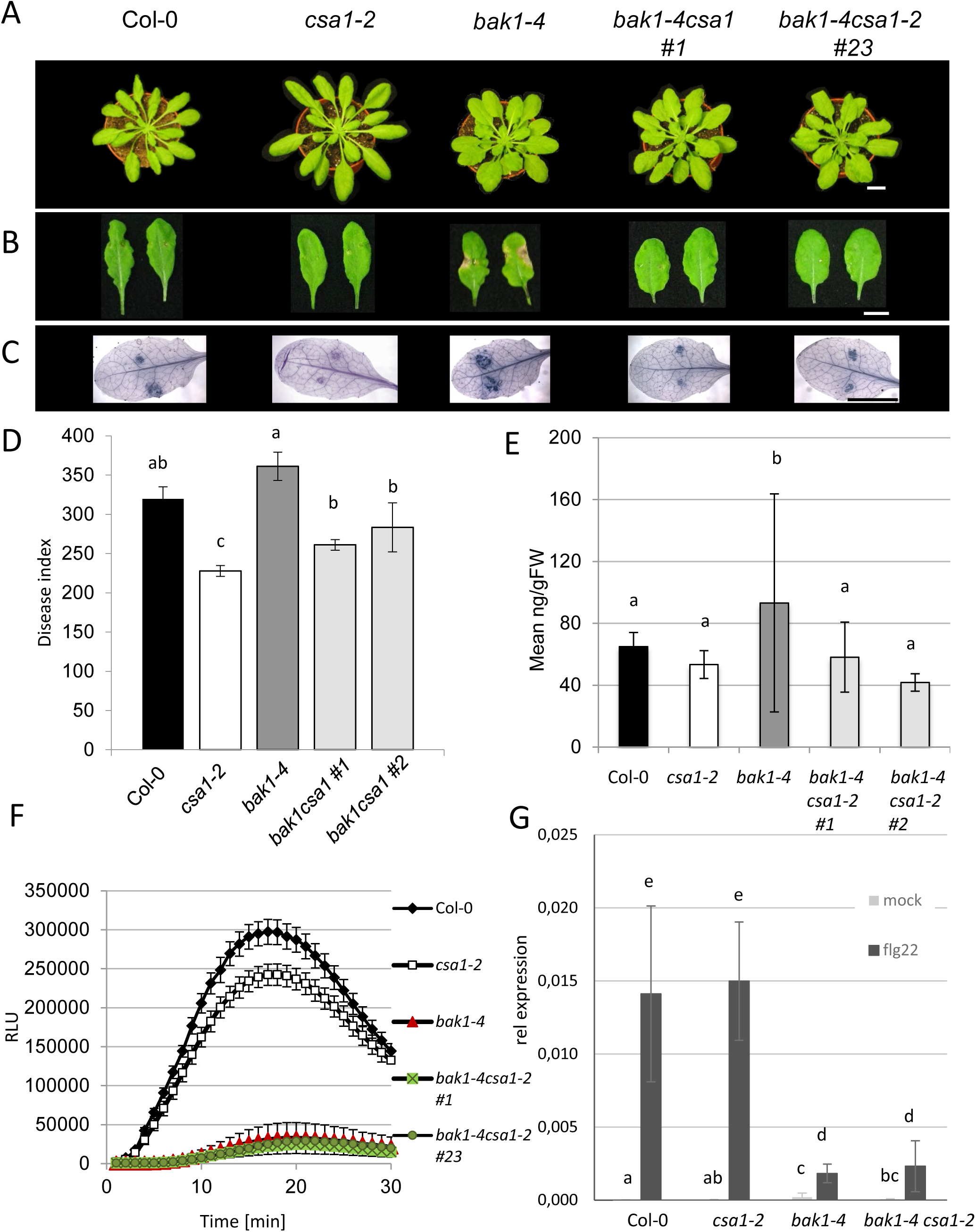
CSA1 is necessary for *bak1*-mediated cell death but not for flg22 responses. **A)** Representative pictures of the morphological phenotype of 6-week-old plants of the indicated genotypes are shown. **(B)** *Alternaria brassicicola* infected leaves of the genotypes shown in (A) 13 days after inoculation. **(C)** Infected leaves of the genotypes shown in (A) stained with trypan blue for cell death. The scale bar in (A), (B) and (C) represents 10 mm. **(D)** Disease indices of *Alternaria brassicicola* infected leaves of the indicated genotypes 13 days after infection shown as mean ± SE (n=12). **(E)** Total salicylic acid levels measured in untreated leaf material of plants of the indicated genotypes relative to the fresh weight. **(F)** ROS production was measured as relative light units (RLU) in a luminol based assay. Leaf pieces of plants of the indicated genotypes were elicited with 100 nM flg22 and ROS production was measured over a period of 30 min. Values are mean ± SE (n=9). **(G)** *FRK1* marker gene expression in leaves of plants of the indicated genotypes was measured by qRT-PCR analysis 3 hours after 100 nM flg22 or mock treatment. *FRK1* expression was normalized to *EF1*α. Results are mean ± SE (n=9). Different letters indicate significant differences according to one-way ANOVA and Tukey’s HSD test (p<0.05). The experiments were repeated at least three times with similar results.

This demonstrates that CSA1 mediates *bak1*-mediated cell death responses and provides evidence for a model with BIR3 being a direct interactor of CSA1 and BAK1, in which CSA1 monitors (“guards”) functional integrity of the BAK1 BIR protein complex that is required for preventing autoimmune cell death in wt plants.

## Discussion

Cell death in *bak1* mutants is a phenomenon that was reported first in 2007 (He et al., 2007; Heese et al., 2007; Kemmerling et al., 2007). Loss-of-function mutants in BAK1-interacting receptors BIR1, BIR2 and BIR3 phenocopy these autoimmune cell death phenotypes (Gao et al., 2009; Halter et al., 2014; Imkampe et al., 2017). Since then, multiple components that are necessary for the execution of the *bak1*-mediated cell death have been revealed. In suppressor screens, often the double mutant *bak1-3 bkk1* and *bak1-4 bkk1* were used as the loss of the two closely related SERK family members results in a strong autoimmune cell death phenotype associated with severe dwarfism or even seedlings lethality (He et al., 2007; Albrecht et al., 2008; Schwessinger et al., 2011; de Oliveira et al., 2016; Du et al., 2016; Gao et al., 2017; Wu et al., 2020). The following components have been shown to suppress BAK1 or BIR cell death: ER quality control components such as ENDOPLASMIC RETICULUM DNAJ DOMAIN-CONTAINING PROTEIN 3B (ERdj3b) and STROMAL CELL-DERIVED FACTOR 2 (SDF2) (Sun et al., 2014), the protein glycosylation machinery component STAUROSPORIN AND TEMPERATURE SENSITIVE3 (STT3a) protein (de Oliveira et al., 2016), the cyclic nucleotide gated ion channel 20 (CNGC20) (Yu et al., 2019), the nucleocytoplasmic trafficking component SUPPRESSOR OF BAK1 BKK1/ NUCLEOPORIN 85 (SBB1/NUP85) and the DEAD-box RNA helicase 1 (DRH1) (Du et al., 2016) and the helper NLR family ADR1 (Wu et al., 2020). A direct detection system or downstream component that senses the absence of BAK1 or BIR proteins was not found.

Effects of *bir1* mutations are partially suppressed by mutations in classical ETI components such as *eds1* and *pad4* mutations and *bak1* cell death is affected by *salicylic acid induction deficient* 2 (*sid2)*, *eds5* mutations, indicating that *bak1*- and *bir1*-mediated cell death is conveyed by SA and potentially involve TNL proteins (Gao et al., 2009; Gao et al., 2017). We also tested multiple ETI downstream components and found that *eds1 (pad4 and sag101)* are partially required for *bak1 bir3* phenotypes, suggesting that TNLs could also be important for this type of cell death. Crosses with the NRG1 family mutants revealed a contribution of NRG1 helper NLRs in *bak1 bir3*-mediated phenotypes. Despite the fact that *bak1 bkk1* cell death is fully dependent on the other helper-NLR family of ADR1 proteins, this shows that *bak1 bir3* mediates cell death is distinct from the *bak1 bkk1* cell death and requires the NRG1s that are linked to TNLs and especially to the CSA1 CHS3 pair of NLRs (Wu et al. 2020; Saile et al, 2020; Wu 2019). If the ADR1 family is additionally involved in *bak1 bir3* cell death needs to be tested in the future. As none of the previously performed genetic screens revealed the direct guard of BAK1 or BIR3, we performed IP-MS analyses with BIR3-YFP and identified the TNL CSA1 as a direct interactor of BIR3 but not BAK1. This explains also why previous interaction studies with BAK1 have not revealed this guard as CSA1 does not directly interact with BAK1 but with its “bodyguard” protein BIR3 and also other BIR proteins.

The IP-MS interactome of BIR3 revealed about 30 RKs as candidate interactors. This broadens the spectrum of RKs that can be assessed by BIR3 and will allow the functional determination of more orphan RKs (Hohmann et al., 2018; Hohmann et al., 2020). RKs associated with BIR proteins are potentially also guarded by NLRs. This might explain how other RKs confer autoimmune cell death such as e.g. SOBIR1 (Gao et al., 2009; Liebrand et al., 2014).

CSA1 and CHS3 are working as a pair in controlling autoimmune cell death (Xu et al., 2015; Castel et al., 2019; Wu et al., 2019). CSA1 can directly interact with BIR3 but not with BAK1, while CHS3 could not be shown to directly interact with BIR3 or BAK1. This suggests a complex with BAK1 interacting strongly with BIR3, BIR3 interacting with CSA1 and CSA1 interacting with CHS3 (Parkes, 2020). The indirect interaction assayed by co-IPs reveals the complex composition also for more distantly associated components. As CSA1 is also necessary for cell death initiated in *bak1* single mutants we propose a model in which CSA1 guards BIR3 directly and the BAK1 BIR3 complex integrity. The activation mechanism of the NLR CSA1 by the absence of BIR3 and/or BAK1 is still elusive, but conformational changes by losing the interaction interface with the receptor complex or transphosphorylation leading to de-repression of the NLR, similar to what is described for RRS1 release and reorganization of the RPS4 complex, appear possible (Williams et al., 2014; Lolle et al., 2020).

Lack of CSA1 can strongly suppress the *bak1 bir3* cell death phenotype, showing that CSA1 is necessary for this cell death execution. Mutations in *CHS3* are less effective and therefore *bak1 bir3*-mediated cell death is different from the *chs3-2D* autoimmune phenotype that is fully dependent on CSA1 (Xu et al., 2015).This is in line with the fact that CSA1 can directly interact with BIR3, but CHS3 does not, and supports the model that CSA1 guards BIR3 directly, potentially bypassing the sensor NLR CHS3 that is only marginally involved in the *bir3 bak1-*mediated cell death.

The phenotype of the triple *bir3 bak1 csa1* mutant does not fully rescue the cell death phenotype exerted by *bak1 bir3* mutations. Other components of the pathway as SA or EDS1 have even less effects. Redundancy within the NLR family might be the reason for this as RPS4 for example is able to complement the reported *csa1* phenotypes (Faigon-Soverna et al., 2006). Also, differential requirement for the two groups of helper NLRs, ADR1s and NRG1s can explain these phenotypes, as BAK1 might preferentially contain cell death via ADR1 suppression, while BIR3 via CSA1 might suppress cytoplasmic cell death activation via the NRG1 family (Wu et al., 2019; Wu et al., 2020).

Though CSA1 is reported to be necessary for *chs3-2D* activated autoimmune cell death, no effector has been identified yet that targets CSA1 or CHS3. BAK1 is targeted by multiple effectors as reported previously for e.g. AvrPto, AvrPtoB, HopF2 and HopB1 (Shan et al., 2008; Zhou et al., 2014; Li et al., 2016). The CSA1 guard has potentially evolved to surveil BAK1 BIR complex integrity to prevent downregulation of PTI by effectors disturbing BAK1 function. Another option is that BAK1-BIR3 complexes are sensed by CSA1 to avoid incompatibility or prevent loss of this central and multifunctional co-receptor complex. A similar scenario was reported for STRUBBELIG RECEPTOR FAMILY 3 (SRF3), an LRR-RLK structurally similar to BIR proteins (Imkampe et al., 2017), that confers hybrid incompatibility along with the R protein RPP1, with SRF3 being guarded by RPP1 to control incompatibility in the absence of pathogens (Alcazar et al., 2010).

BAK1 as a classical PTI co-receptor is linked to an NLR that is classically part of the ETI pathway. Recent work suggests that PTI is required for proper ETI and HR cell death, but the mechanism of this interplay is still elusive (Ngou et al., 2021; Yuan et al., 2021). Mutants in *CSA1* are slightly more susceptible to bacterial pathogens (Faigon-Soverna et al., 2006). CSA1 can interact with components from both extracellular and intracellular immune receptors and may provide a link between both immune systems. Early PTI responses are not altered in *csa1* mutants (Figure 6, 7) but bacterial resistance to *Pto* DC3000 avrRpt2 is impaired, indicating that plant resistance is affected by loss of *CSA1* (Faigon-Soverna et al., 2006). This points to an effective interaction of the NLR protein CSA1 with pattern recognition receptor complexes, and lack of interaction result in HR cell death, potentially enhancing plant resistance to biotrophic pathogens (Faigon-Soverna et al., 2006). Depletion of BAK1 after infection might be used by fungal pathogens to improve virulence by cell death induction in the plant (Yamada et al., 2016).

## Conclusion

Here we describe the TNL protein CSA1 to interact physically and genetically with BIR3 and thereby senses BIR3/BAK1-complex integrity and induces cell death when the complex integrity is disturbed. The identification of CSA1 helps explaining the cell death phenotypes of *bak1* and *bir3* mutants as a result of a surveillance system that controls the integrity of this essential PTI co-receptor complex by an NLR protein typically involved in ETI signaling. To find out if this involves microbial effectors that target the PTI co-receptors or if activation of BAK1 results in additional activation of ETI-responses that amplify PTI responses or if it is a result of linking NLR-mediated autoimmune cell death with PTI responses will be an interesting challenge for the future.

## Materials and Methods

### Plant material and growth conditions

Plants were grown for 5 to 6 weeks on soil in growth chambers under short day conditions (8 hr light, 16 hr dark; 22°C; 110 µEm^−2^ s^−1^), for five to six weeks under long day conditions (16hr light, 8 hr dark) or on ½ MS medium.

Mutants used are *bir3-2* (Salk_116632), *bak1-4* (SALK_116202), *csa1-2* (SALK_023219) *chs3-3* (SALK_063886), *eds1-12 (Ordon et al., 2017)*, *pad 4-1 (Glazebrook et al., 1996), sag101-2 (Feys et al., 2005)*, *nrg1.1, nrg1.2*. (Castel et al., 2019). Lines were crossed with *bak1-4* and *bak1-4 bir3-2.* NahG expressing lines were crossed with *bak1-4* and *bak1-4 bir3-2* lines. All lines were genotyped with the primers indicated in STable2.

Constructs for stable transformed Arabidopsis and for transient expression in *N. benthamiana* pBIR3-BIR3-eGFP, pBAK1-BAK1-eGFP, 35S-BIR2-YFP, 35S-BIR1-eGFP were described previously (Halter et al., 2014; Imkampe et al., 2017). To clone the genomic fragment containing *CSA1* (*At5g178880*), TAC clone JAtY79I19 obtained from Arabidopsis accession Col-0 was digested with *Kpn*I (NEB) and *Sal*I (NEB), and a 7,959 bp fragment harbouring *proCSA1:gCSA1* was purified from agarose gel and cloned into *Kpn*I and *Sal*I digested *pCambia2300*.

Cloning of *35S-CSA1-V5* and *35S-CHS3-V5* was carried out using Uracil-Specific Excision Reagent (USER) method (Nour-Eldin et al., 2010). *CSA1* and *CHS3* were amplified from TAC clone JAtY79I19 with the primers USER-CSA1-F/USER-CSA1-R and USER-CHS3-F/USER-CHS3-R, respectively. The resulting PCR products together with V5-tag amplicons (amplified with USER-V5-F/USER-V5-R) were cloned into *LBJJ234* vector pre-linearized with *Pac*I and Nt.*Bbvc*I as described by (Redkar et al., 2021).

GFP-fusion proteins of CSA1 and CHS3 were generated by amplification from Col-0 cDNA with the primers listed in STable 2 and cloning into *pENTR-TOPO* (Thermofischer) and recombination into *pB7FWG2* (Karimi et al., 2002).

Genomic clones for complementation were amplified from *pCAMBIA2300* clones described above with the primers listed in STable2, cloned in *pENTR-TOPO* (Thermofischer) and recombined in *pFAST-G01.* The constructs were transformed in *Agrobacterium* strain GV3101 and transformed in *bak1 csa1* and *bak1 bir3 csa1* plants by floral dipping (Clough and Bent, 1998).

### Infection procedures

*Alternaria brassicicola* infection assays were carried out as described by Kemmerling et al. (2007).

### Histochemical assays

Cell death and fungal mycelium was detected with trypan blue staining as described in Kemmerling et al. (2007).

### Oxidative burst measurements

Oxidative burst was measured with a luminol-based assay as described in Halter et al. (2014)

### RT-PCR analysis

Transcript levels were analyzed by quantitative RT-PCR (qRT-PCR) as described by Mosher et al. (2013) with primers listed in Supplemental Table S2.

### Hormone measurements

Salicylate contents were measured as described by Lenz et al. (2011).

### Transient expression in *Nicotiana benthamiana*

BIR3, BAK1, BIR2, BIR1, CSA1 CHS3 constructs described above were transformed into Agrobacterium strain GV3101 or C58 and transiently expressed in *N. benthamiana* as described in Halter et al. (2014).

### Co-immunoprecipitations

Leaves were ground in liquid nitrogen, and 250 µl extraction buffer (50 mM Tris-HCl pH 8.0, 150 mM NaCl, 1 % Nonidet P40, proteinase inhibitor cocktail (Roche)) per 200 mg tissue powder was added. Samples were homogenized and incubated for 1 h at 4°C under gentle shaking. Samples were centrifuged two times at 4°C and 14,000 rpm for 10 min to obtain a clear protein extract. After washing with extraction buffer, either 15 µl of GFP-trap or V5-trap beads (Sigma, Chromotec) were used. Supernatants containing equal amounts of protein were incubated for 1 h at 4°C with the beads. Beads were washed two times with 50 mM Tris-HCl pH 8.0, 150 mM NaCl and one time with 50 mM Tris-HCl pH 8.0, 50 mM NaCl before adding SDS sample buffer and heating at 95°C for 5 min.

### SDS-PAGE and immunoblotting

Proteins were separated, blotted, and incubated with antibodies as described by Schulze et al. (2010) but using 8% SDS gels and the following antibody dilutions: anti-GFP (Abcam), 1:3,000; anti-HA (Sigma), 1:2,000; anti-V5 (Sigma), 1:2,000; anti-BAK1 (Agrisera), 1:3,000; anti-BIR3 antibodies were obtained from rabbits immunized with the peptide CVGSRDSNDSSFNN fused to KLH (Agrisera) 1:500; anti-luciferase (Sigma), 1:3,000; anti-VP16 (Santa Cruz), 1:1000; anti-ATPase (Agrisera), 1:5,000; 1:5,000; anti-rabbit (Sigma),1:50,000; anti-goat (Sigma), 1:10,000; anti-mouse (Sigma), 1:10,000. Chemiluminescence was detected with the ECL Western blotting detection system (GE Healthcare) and a CCD camera (Amersham Imager 600). If figures were reconstituted from images of blots, lanes from the same blot are shown in one panel of a figure even if they were in a different order on the original blot (separated by dotted lines). Data from different blots are shown in separate figure parts.

### Mass spectrometry analyses

Coomassie-stained gel pieces were digested in gel with trypsin as described previously (Borchert et al., 2010). After desalting using C18 Stage tips (Rappsilber et al., 2007), extracted peptides were separated on an EasyLC nano-HPLC coupled to a Q Exactive HF mass spectrometer (both Thermo Fisher Scientific) as described elsewhere (Kelstrup et al., 2014) with slight modifications: the peptide mixtures were injected onto the column in HPLC solvent A (0.1% formic acid) at a flow rate of 500 nl/min and subsequently eluted with a 57minute segmented gradient of 10–33-50-90% of HPLC solvent B (80% acetonitrile in 0.1% formic acid) at a flow rate of 200 nl/min. Full scan was acquired in the mass range from m/z 300 to 1650 at a resolution of 120,000 followed by HCD fragmentation of the 7 most intense precursor ions. High-resolution HCD MS/MS spectra were acquired with a resolution of 60,000. The target values for the MS scan and MS/MS fragmentation were 3x 10^6^ and 10^5^ charges, respectively. Precursor ions were excluded from sequencing for 30 s after MS/MS.

### Data processing

Acquired MS spectra were processed with MaxQuant software package version 1.5.1.0 (Cox and Mann, 2008) with integrated Andromeda search engine (Cox et al., 2011). The database search was performed against a target-decoy *Arabidopsis thaliana* database obtained from Uniprot, containing 33,351 protein entries, and 245 commonly observed contaminants. Endoprotease trypsin was defined as protease with a maximum of two missed cleavages. Oxidation of methionine and N-terminal acetylation were specified as variable modifications, whereas carbamidomethylation on cysteine was set as a fixed modification. Initial maximum allowed mass tolerance was set to 4.5 ppm (for the survey scan) and 20 ppm for HCD fragment ions. Peptide, protein and modification site identifications were reported at a false discovery rate (FDR) of 0.01, estimated by the target/decoy approach (Elias and Gygi, 2007).

### Yeast split ubiquitin assay

BAK1, BIR3, CSA1 and CHS3 cDNAs were amplified with the primers listed in STable 2 and cloned into pCR8 according to the manufacturers protocol and recombined into pXNubA22-Dest and pMetYC-Dest vectors for direct interaction assays of membrane proteins in yeast using the split-ubiquitin system (SUS) (Grefen et al., 2009). Yeast assays were performed as described in Halter et al., (2014)

### Split-Luciferase assay

BIR3 and BAK1 cDNA and CSA1 and CHS3 genomic coding sequences were amplified with the primers listed in STable 2 and classically cloned into the KpnI SalI restriction sites of the vectors pCAMBIA-NLuc and pCAMBIA-CLuc for expression in *Nicotiana benthamiana*. The Split-Luciferase assays were performed as described in Zhou et al. (2018).

### Statistical Methods

Statistical significance between two samples was tested with Student’s t-test, between groups statistical significance was analyzed using one-way ANOVA combined with Tukey’s honest significant difference (HSD) test. Significant differences are indicated with different letters (p < 0.05).

### Accession numbers

*BIR3: At1g27190; BIR2: At3g28450; BIR1: At3g48380; BAK1/SERK3: At4g33430; BKK1/SERK4: At2g13790; CSA1: AT5G17880; CHS3:At5g17890; EDS1:At3g48090; PAD4: At3g53420; SAG101: At5g14930, NRG1.1: At5g66900 NRG1.2: At5g69910*.

## Acknowledgements

This study was supported by the DFG SFB1101 D03 project to B.K. and DFG SFB1101 D09 project to F.E.K. We thank Silke Wahl and Irina Droste-Borel for technical support for the proteomics assays, J.-M. Zhou for providing Split-Luc vectors and Thorsten Nürnberger for critical reading of the manuscript.

## Author Contributions

**SS** and **LY** designed research, performed research and analyzed data; **AE, DK, MFW and MS** performed research and analyzed data; **SCS** performed research, analyzed data and designed research; **LL** provided materials for this research; **FEK** analyzed data, designed research and provided materials for this research, **VC** designed research and provided materials for this research and **BK** designed research, analyzed data and wrote the manuscript. All authors revised the manuscript.

## Competing Interests statement

The authors declare no competing interests.

## Supplemental Tables

Supplemental Table1: BIR3 interacting RKs.xls

Supplemental Table 2: Primers used in this study

## Supplemental Figure titles

Supplemental Figure1. Reduced SA levels by *NahG* expression and mutation in *PAD4* and *SAG101* can weakly suppress the dwarf phenotype of *bak1 bir3* mutants

Supplemental Figure 2: Helper NLRs NRG1-1 and NRG1-2 are necessary for *bak1-4bir3-2* double mutant phenotype

Supplemental Figure 3: Mass spectrometric identification of the BIR3 interacting protein CSA1

Supplemental Figure 4: CSA1 can interact with BIR1 and BIR2

Supplemental Figure 5: CSA1 localizes preferentially to microsomal fractions

Supplemental Figure 6: Expression controls for Split-luciferase assays

Supplemental Figure 7: Expression controls for Split-ubiquitin assays

Supplemental Figure 8: Sequence and domain structure of CHS3

Supplemental Figure 9: CHS3 can interact with BIR1 and BIR2

Supplemental Figure 10: CSA1 is not sufficient to suppress *bir1*-mediated dwarfism

Supplemental Figure 11. Mutation in *chs3* partially suppresses cell death phenotypes in *bak1 bir3* double mutants

Supplemental Figure 12: Expression of CSA1 can complement the *bak1 bir3 csa1* triple mutant phenotype

Supplemental Figure 13: Expression of CSA1 can complement the *bak1 csa1* double mutant phenotype

## Notes

### Competing Interest Statement

The authors have declared no competing interest.

