## Supplemental Figures 1-13, Supplemental Table 2 for "The TIR-NBS-LRR protein CSA1 is required for autoimmune cell death in Arabidopsis pattern recognition co-receptor *bak1* and *bir3* mutants"

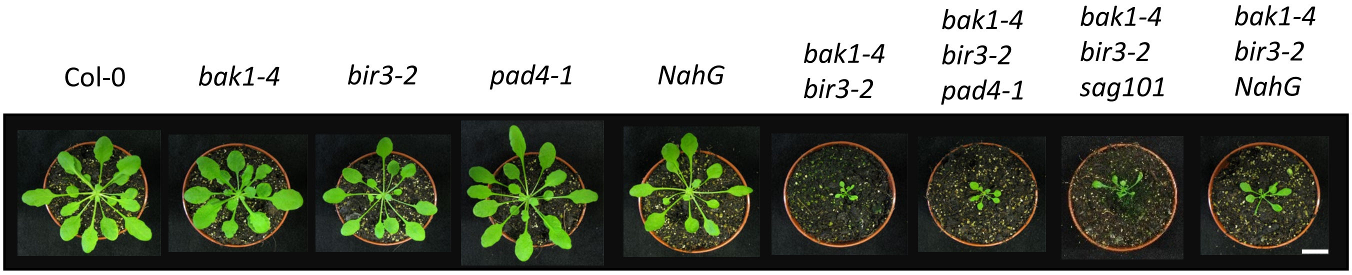


**Supplemental Figure1. Reduced SA levels by *NahG* expression and mutation in *PAD4* and *SAG101* can weakly suppress the dwarf phenotype of *bak1 bir3* mutants**
Representative pictures of 6-week-old plants of the indicated genotypes are shown. The scale bar represents 1 cm.

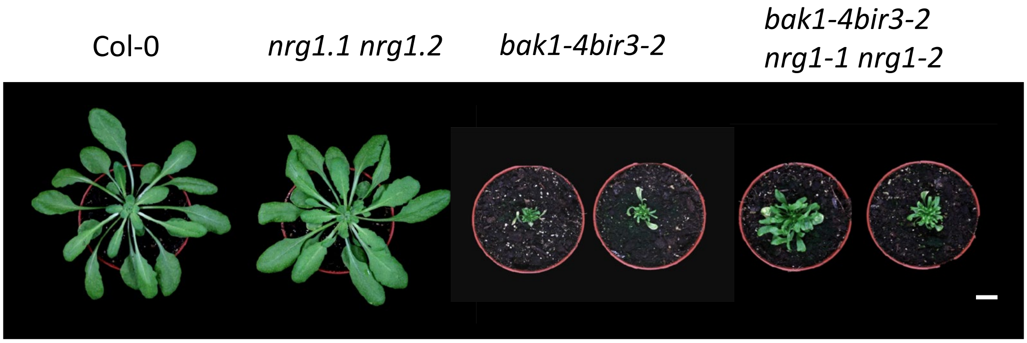


**Supplemental Figure 2: Helper NLRs NRG1-1 and NRG1-2 are necessary for *bak1-4bir3-2* double mutant phenotypes**

**(A)** Representative pictures of the morphological phenotype of 6-week-old Col-0, *nrg1-1 nrg1-2* and *bir3-2 bak1-4* double mutant and the quadruple mutant *bak1-4 bir3-2 nrg1-1 nrg1-2.* The scale bar represents 1 cm.

A

| Raw file | Scan | Method | Score | m/z | Gene names |
|---|---|---|---|---|---|
| 20150508_CO_0552BiKe_R02_01 | 7308 | FTMS; HCD | 76.22 | 485.78 | CSA1 |

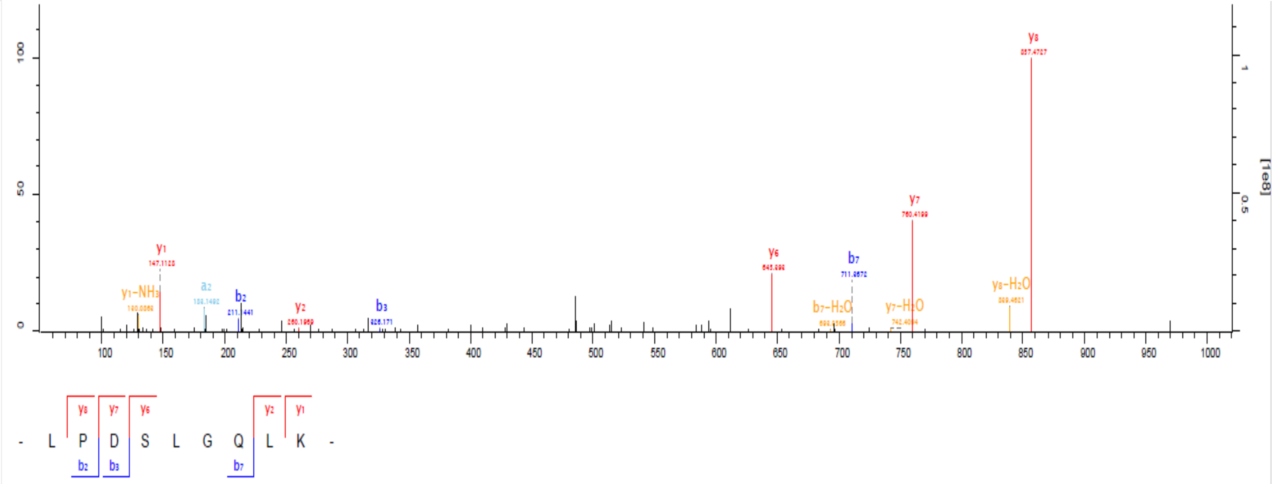


B

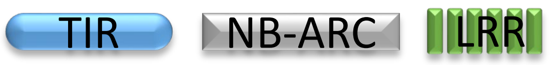


CSA1 At5g17880

C

```
   1 MTSSSSWVKT DGETPQDQVF INFRGVELRK NFVSHLEKGL KRKGINAFID
  51 TDEEMGQELS VLLERIEGSR IALAIFSPRY TESKWCLKEL AKMKERTEQK
 101 ELVVIPIFYK VQPVTVKELK GDFGDKFREL VKSTDKKTKK EWKEALQYVP
 151 FLTGIVLDEK SDEDEVINII IRKVKEILNR RSEGPPSKCS ALPPQRHQKR
 201 HETFWGIELR IKQLEEKLRF GSDETTRTIG VVGMPGIGKT TLATMLYEKW
 251 NDRFLRHVLI RDIHEASEED GLNYLATKFL QGLLKVENAN IESVQAAHEA
 301 YKDQLLETKV LVILDNVSNK DQVDALLGER NWIKKGSKIL ITTSDKSLMI
 351 QSLVNDTYEV PPLSDKDAIK HFIRYAFDGN EGAAPGPGQG NFPKLSKDFV
 401 HYTKGNPLAL QMLGKELLGK DESHWGLKLN ALDQHHNSPP GQSICKMLQR
 451 VWEGSYKALS QKEKDALLDI ACFRSQDENY VASLLDSDGP SNILEDLVNK
 501 FMINIYAGKV DMHDTLYMLS KELGREATAT DRKGRHRLWH HHTIIAVLDK
 551 NKGGSNIRSI FLDLSDITRK WCFYRHAFAM MRDLRYLKIY STHCPQECES
 601 DIKLNFPEGL LLPLNEVRYL HWLKFPLKEV PQDFNPGNLV DLKLPYSEIE
 651 RVWEDNKDAP KLKWVNLNHS KKLNTLAGLG KAQNLQELNL EGCTALKEMH
 701 VDMENMKFLV FLNLRGCTSL KSLPEIQLIS LKTLILSGCS KFKTFQVISD
 751 KLEALYLDGT AIKELPCDIG RLQRLVMLNM KGCKKLKRLP DSLGQLKALE
 801 ELILSGCSKL NEFPETWGNM SRLEILLLDE TAIKDMPKIL SVRRLCLNKN
 851 EKISRLPDLL NKFSQLQWLH LKYCKNLTHV PQLPPNLQYL NVHGCSSLKT
 901 VAKPLVCSIP MKHVNSSFIF TNCNELEQAA KEEIVVYAER KCHLLASALK
 951 RCDESCVPEI LFCTSFPGCE MPSWFSHDAI GSMVEFELPP HWNHNRLSGI
1001 ALCVVVSFKN CKSHANLIVK FSCEQNNGEG SSSSITWKVG SLIEQDNQEE
1051 TVESDHVFIG YTNCLDFIKL VKGQGGPKCA PTKASLEFSV RTGTGGEATL
1101 EVLKSGFSFV FEPEENRVPS PRNDDVKGKV KINKTPSANG CFKDQAKGNE
1151 SPKGQWQTYI ENSSTNIPSE AHSSQKTGFN GFNGMYSVCV LYEMYSH
```

**Supplemental Figure 3: Mass spectrum of the BIR3-interacting protein CSA1**

**A)** Product ion spectrum (b and y ions) of the CSA1 peptide generated from a tryptic digest of a BIR3-eGFP IP of whole Arabidopsis seedlings using ion trap LC/MS/MS analysis. The spectrum was verified by manual inspection. **B)** The identified peptide belongs to the NLR protein CSA1, a protein containing the following domains: Toll-Interleukin receptor (TIR), nucleotide binding APAF-1 (apoptotic protease-activating factor-1), R proteins and CED-4 (*Caenorhabditis elegans* death-4 protein) (NB-ARC) and the leucine-rich repeat (LRR) domain. **C)** Sequence of CSA1 with the domains identified with Interpro Scan labelled with the following color code: TIR domain (blue), NB-ARC-domain (grey) including the P-loop (underlined), LRR domain (green). The peptide identified in the Co-IPs by MS analyses is labelled in red.

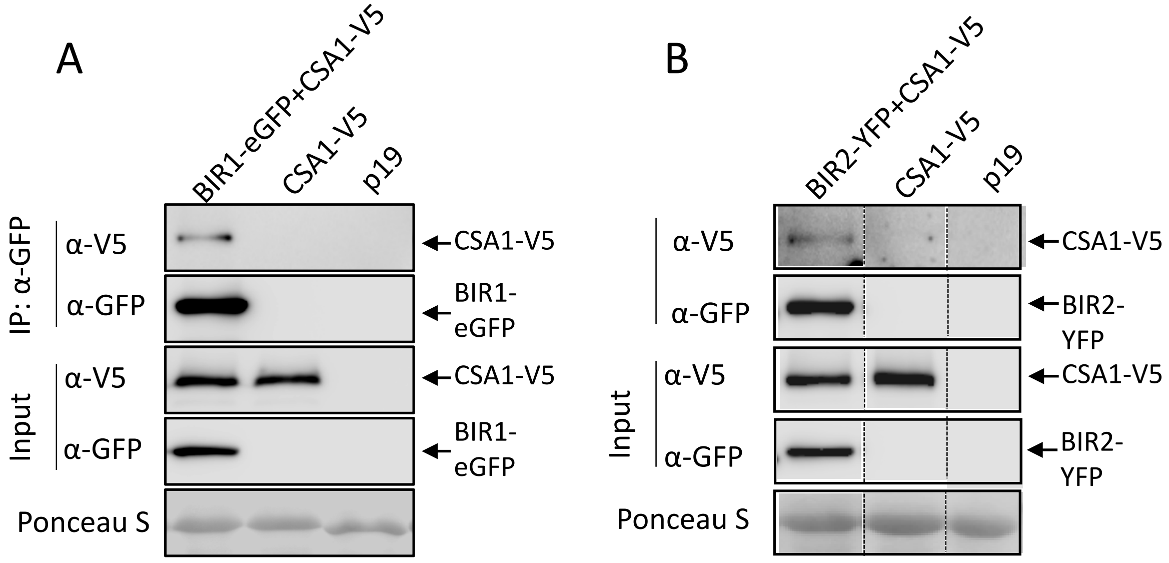


**Supplemental Figure 4: CSA1 can interact with BIR1 and BIR2**

Western Blots after co-immunoprecipitation with GFP-traps of transiently in *Nicotiana benthamiana* expressed **A)** BIR1-eGFP and **B)** BIR2-YFP and CSA1-V5 detected with α-GFP and α-V5 antibodies. Protein input is shown by Western blot analysis of protein extracts before IP and antibodies against the respective tags. p19 is a silencing inhibitor expressed alone as a background control. Ponceau S staining shows protein loading. Dotted lines indicate cut and rearranged parts of the same blot.

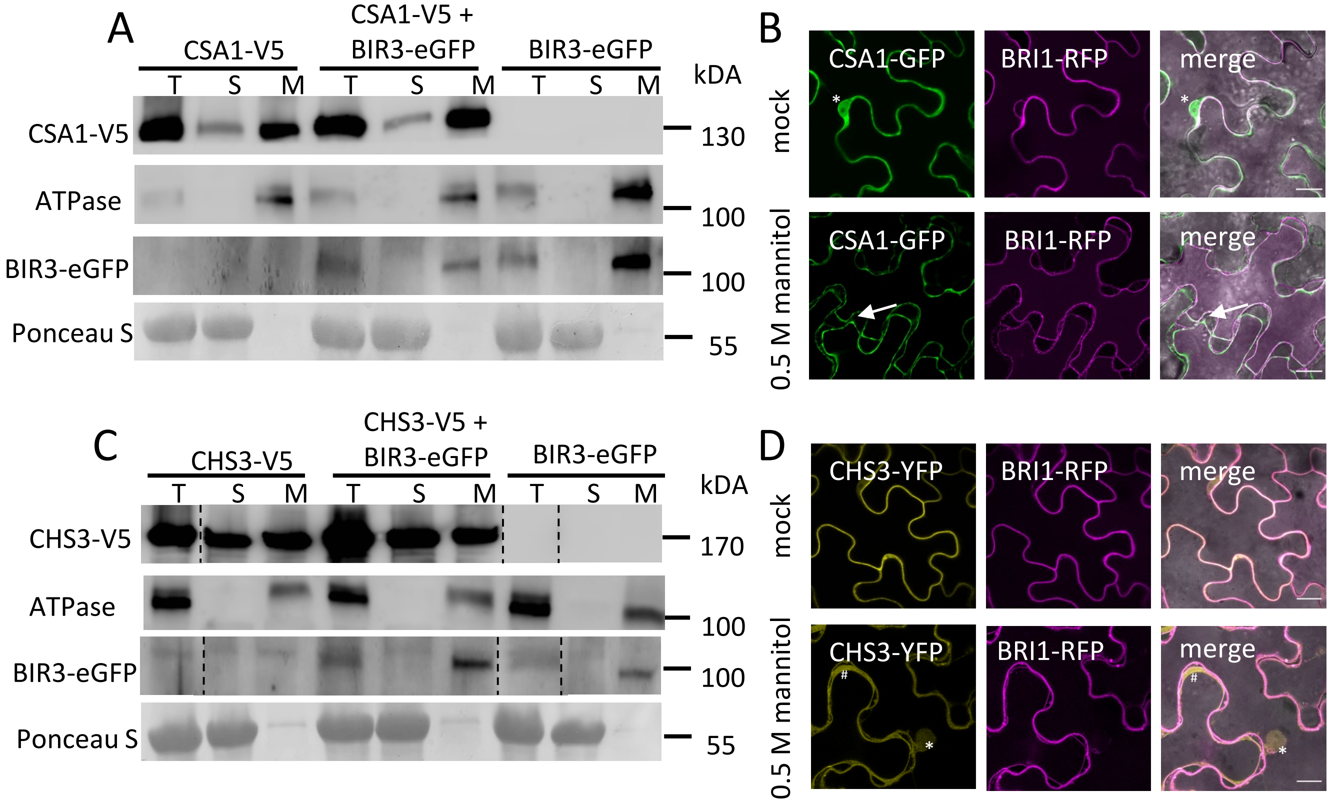


**Supplemental Figure 5: CSA1 localizes preferentially to microsomal fractions**

**A)** CSA1-V5, **C)** CHS3-V5 and BIR3-eGFP were transiently expressed alone and in combination in *Nicotiana benthamiana* and total (T), soluble (S) and microsomal (M) protein fractions were analyzed by Western blot and detected with α-V5, α-ATPase and α-GFP antibodies to detect **A)** CSA1 and **C)** CHS3 protein in the different fractions. An ATPase and BIR3-eGFP are shown as markers for membrane localization. Ponceau S staining shows protein loading.

Confocal laser scanning microscopy images of **B)** CSA1-GFP and **D)** CHS3-GFP transiently expressed in *Nicotiana benthamiana* and BRI1-RFP as a membrane localized control. 0,5M mannitol was used to induce plasmolysis. Merged figures show co-localization of CSA1 or CHS3 and BRI1 in the red and green channel. The arrows mark Hechtian strands, asterisks the nuclei, and hashtags cytoplasm, size bars represent 20µm.

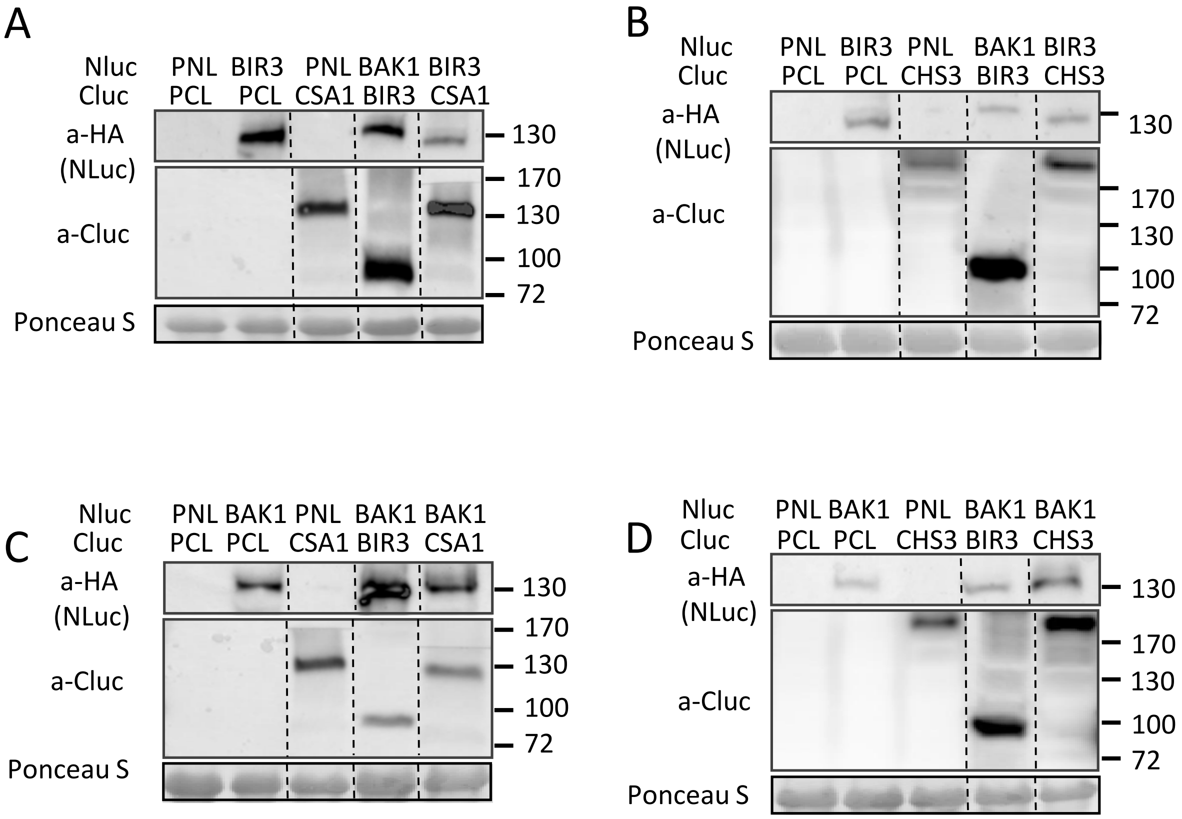


**Supplemental Figure 6: Expression controls for Split-luciferase assays**

Western blots of in *Nicotiana benthamiana* expressed fusion proteins as shown in **A)** Fig. 2, **B)** Fig. 3, **C)** Fig. 4, **D)** Fig. 5 detected with $\alpha$-HA (Nluc) and $\alpha$-luciferase (Cluc) antibodies. Ponceau S staining shows protein loading. Dotted lines indicate cut and rearranged parts of the same blot.

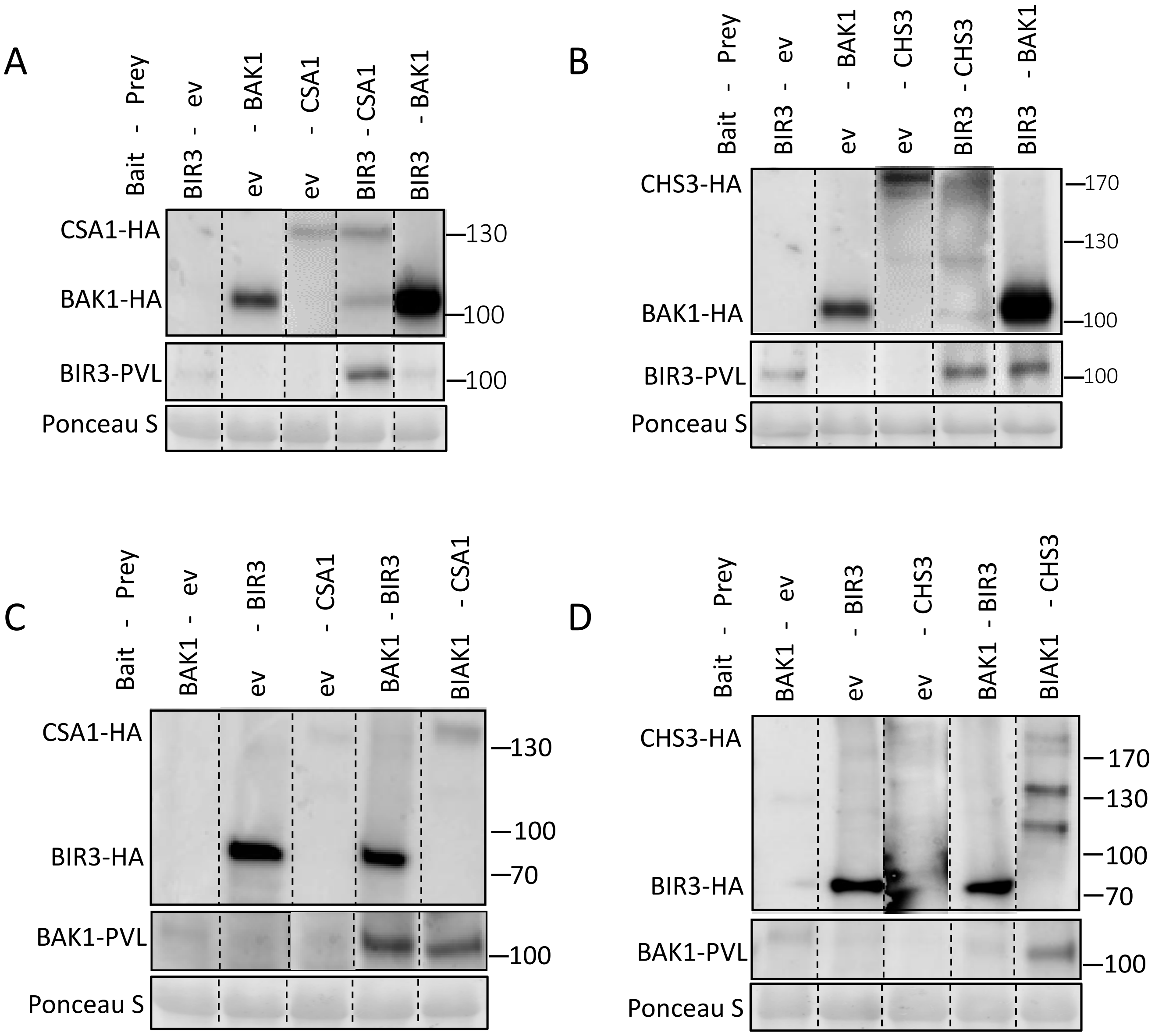


**Supplemental Figure 7: Expression controls for Split-ubiquitin assays**

Western blots of in yeast expressed fusion proteins as shown in **A)** Fig. 2, **B)** Fig. 3, **C)** Fig. 4, **D)** Fig. 5 detected with α-HA (prey) and α-VP16 (bait) antibodies. Ponceau S staining shows protein loading. Dotted lines indicate cut and rearranged parts of the same blot.

A

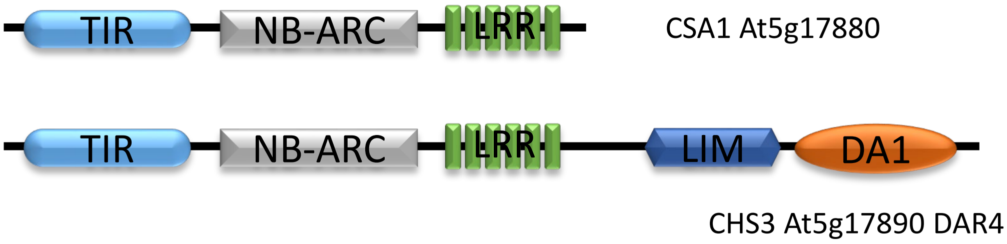


B

```
1    MEPPAARVTP SIKADCSHSV NIICEETVLH SLVSHLSAAL RREGISVFVD
51   ACGLQETKFF SIKQNQPLTD GARVLVVVIS DEVEFYDPWF PKFLKVIQGW
101  QNNGHVVVPV FYGVDSLTRV YGWANSWLEA EKLTSHQSKI LSNNVLTDSE
151  LVEEIVRDVY GKLYPAERVG IYARLLEIEK LLYKQHRDIR SIGIWGMPGI
201  GKTTLAKAVF NHMSTDYDAS CFIENFDEAF HKEGLHRLLK ERIGKILKDE
251  FDIESSYIMR PTLHRDKLYD KRILVVLDDV RDSLAAESFL KRLDWFGSGS
301  LIIITSVDKQ VFAFCQINQI YTVQGLNVHE ALQLFSQSVF GINEPEQNDR
351  KLSMKVIDYV NGNPLALSIY GRELMGKKSE METAFFELKH CPPLKIQDVL
401  KNAYSALSDN EKNIVLDIAF FFKGETVNYV MQLLEESHYF PRLAIDVLVD
451  KCVLTISENT VQMNNLIQDT CQEIFNGEIE TCTRMWEPSR IRYLLEYDEL
501  EGSGETKAMP KSGLVAEHIE SIFLDTSNVK FDVKHDAFKN MFNLKFLKIY
551  NSCSKYISGL NFPKGLDSLP YELRLLHWEN YPLQSLPQDF DFGHLVKLSM
601  PYSQLHKLGT RVKDLVMLKR LILSHSLQLV ECDILIYAQN IELIDLQGCT
651  GLQRFPDTSQ LQNLRVVNLS GCTEIKCFSG VPPNIEELHL QGTRIREIPI
701  FNATHPPKVK LDRKKLWNLL ENFSDVEHID LECVTNLATV TSNNHVMGKL
751  VCLNMKYCSN LRGLPDMVSL ESLKVLYLSG CSELEKIMGF PRNLKKLYVG
801  GTAIRELPQL PNSLEFLNAH GCKHLKSINL DFEQLPRHFI FSNCYRFSSQ
851  VIAEFVEKGL VASLARAKQE ELIKAPEVII CIPMDTRQRS SFRLQAGRNA
901  MTDLVPWMQK PISGFSMSVV VSFQDDYHND VGLRIRCVGT WKTWNNQPDR
951  IVERFFQCWA PTEAPKVVAD HIFVLYDTKM HPSDSEENHI SMWAHEVKFE
1001 FHTVSGENNP LGASCKVTEC GVEVITAATG DTSVSGIIRE SETITIIEKE
1051 DTIIDEEDTP LLSRKPEETN RSRSSSELQK LSSTSSKVRS KGNVFWKWLG
1101 CFPLQPKNLR SRSRRTTALE EALEEALKER EKLEDTRELQ IALIESKKIK
1151 KIKQADERDQ IKHADEREQR KHSKDHEEEE IESNEKEERR HSKDYVIEEL
1201 VLKGKGKRKQ LDDDKADEKE QIKHSKDHVE EEVNPPLSKC KDCKSAIEDG
1251 ISINAYGSVW HPQCFCCLRC REPIAMNEIS DLRGMYHKPC YKELRHPNCY
1301 VCEKKIPRTA EGLKYHEHPF WMETYCPSHD GDGTPKCCSC ERLEHCGTQY
1351 VMLADFRWLC RECMDSAIMD SDECQPLHFE IREFFEGLHM KIEEEFPVYL
1401 VEKNALNKAE KEEKIDKQGD QCLMVVRGIC LSEEQIVTSV SQGVRRMLNK
1451 QILDTVTESQ RVVRKCEVTA ILILYGLPRL LTGYILAHEM MHAYLRLNGY
1501 RNLNMVLEEG LCQVLGYMWL ECQTYVFDTA TIASSSSSSR TPLSTTTSKK
1551 VDPSDFEKRL VNFCKHQIET DESPFFGDGF RKVNKMMASN NHSLKDTLKE
1601 IISISKTPQY SKL
```

**Supplemental Figure 8: Sequence and domain structure of CHS3**

**A)** CHS3 contains Toll-Interleukin receptor (TIR), nucleotide binding APAF-1 (apoptotic protease-activating factor-1), R proteins and CED-4 (*Caenorhabditis elegans* death-4 protein) (NB-ARC) and leucine-rich repeat (LRR) domains plus integrated domains (CC: coiled coil, LIM: LIN-11, Isl-1 and MEC-3 and DA-1: DA-1-like protease domain). **B)** Sequence of CHS3 with predicted domains based on Interpro Scan. Domains are marked with the following color code: TIR domain (blue), NB-ARC-domain (grey) including the P-loop (underlined), LRR-domain (LRR / green), coiled-coil domain (CC / yellow), LIM-domain (LIM / dark blue) and a zinc protease domain including the underlined HExxH motif (orange), required for protease activity.

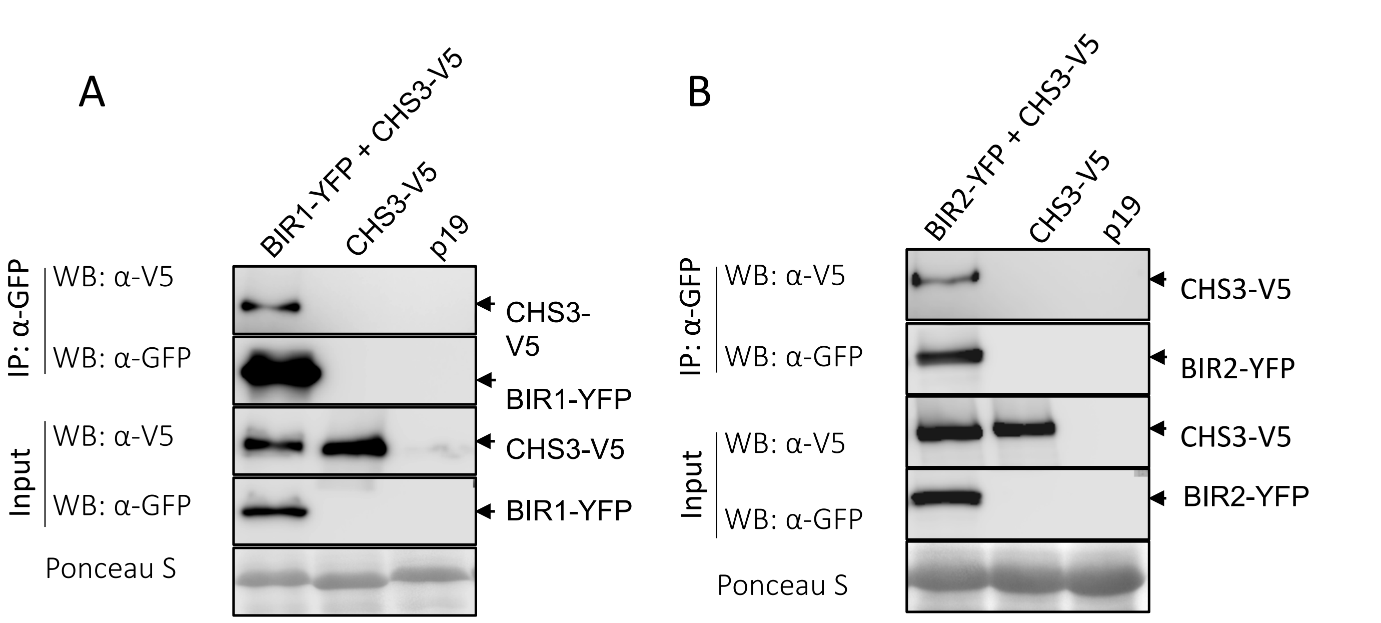


**Supplemental Figure 9: CHS3 can interact with BIR1 and BIR2**

Western Blots after co-immunoprecipitation with GFP-traps of transiently in *Nicotiana benthamiana* expressed **A)** BIR1-eGFP and **B)** BIR2-YFP and CHS3-V5 detected with α-GFP and α-V5 antibodies. Protein input is shown by Western blot analysis of protein extracts before IP and antibodies against the respective tags. p19 is a silencing inhibitor expressed alone as a background control. Ponceau S staining shows protein loading. Dotted lines indicate cut and rearranged parts of the same blot.

Col-0 | *csa1-2* | *bir1-1* | *csa1 bir1-1* #1 | *csa1 bir1-1* #2

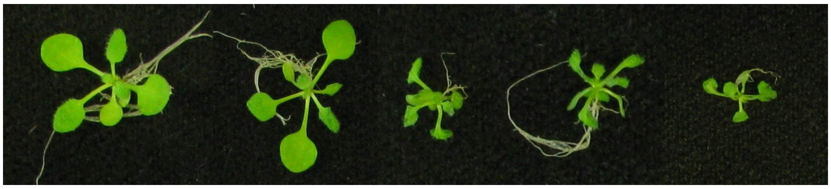

**Supplemental Figure 10: CSA1 is not sufficient to suppress *bir1*-mediated dwarfism**
Representative pictures of two-week old Arabidopsis seedlings of the indicated genotypes are shown.

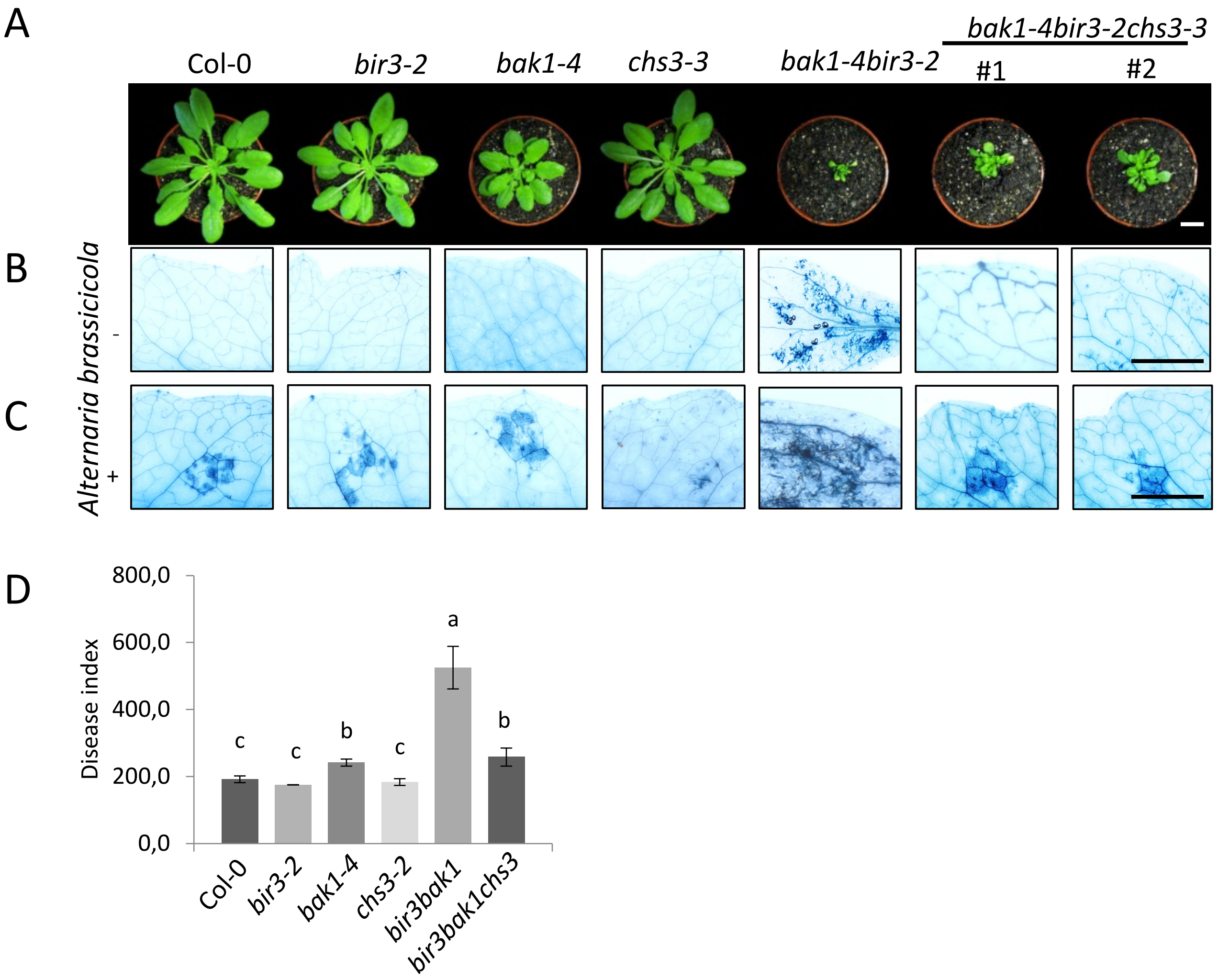


**Supplemental Figure 11. Mutation in *chs3* partially suppresses cell death phenotypes in *bak1 bir3* double mutants**

**(A)** Representative pictures of the morphological phenotype of 6-week-old Col-0, *bir3-2, bak1-4, chs3-2* and the double mutant *bak1-4 bir3-2* and the triple mutant *bak1-4 bir3-2 chs3-2.* The scale bar represents 1 cm. **(B)** Uninfected leaves of the genotypes shown in (A) stained with trypan blue for cell death. **(C)** Leaves of the same genotypes as in (A) and (B) droplet-infected with *Alternaria brassicicola* and trypan blue stained. The scale bar in (A) and (B) represents 5 mm. **(D)** Disease indices of *Alternaria brassicicola* infected leaves of the indicated genotypes 13 days after infection shown as mean ± SE (n=12). Different letters indicate significant differences according to one-way ANOVA and Tukey's HSD test ($p<0.05$). The experiments were repeated at least three times with similar results.

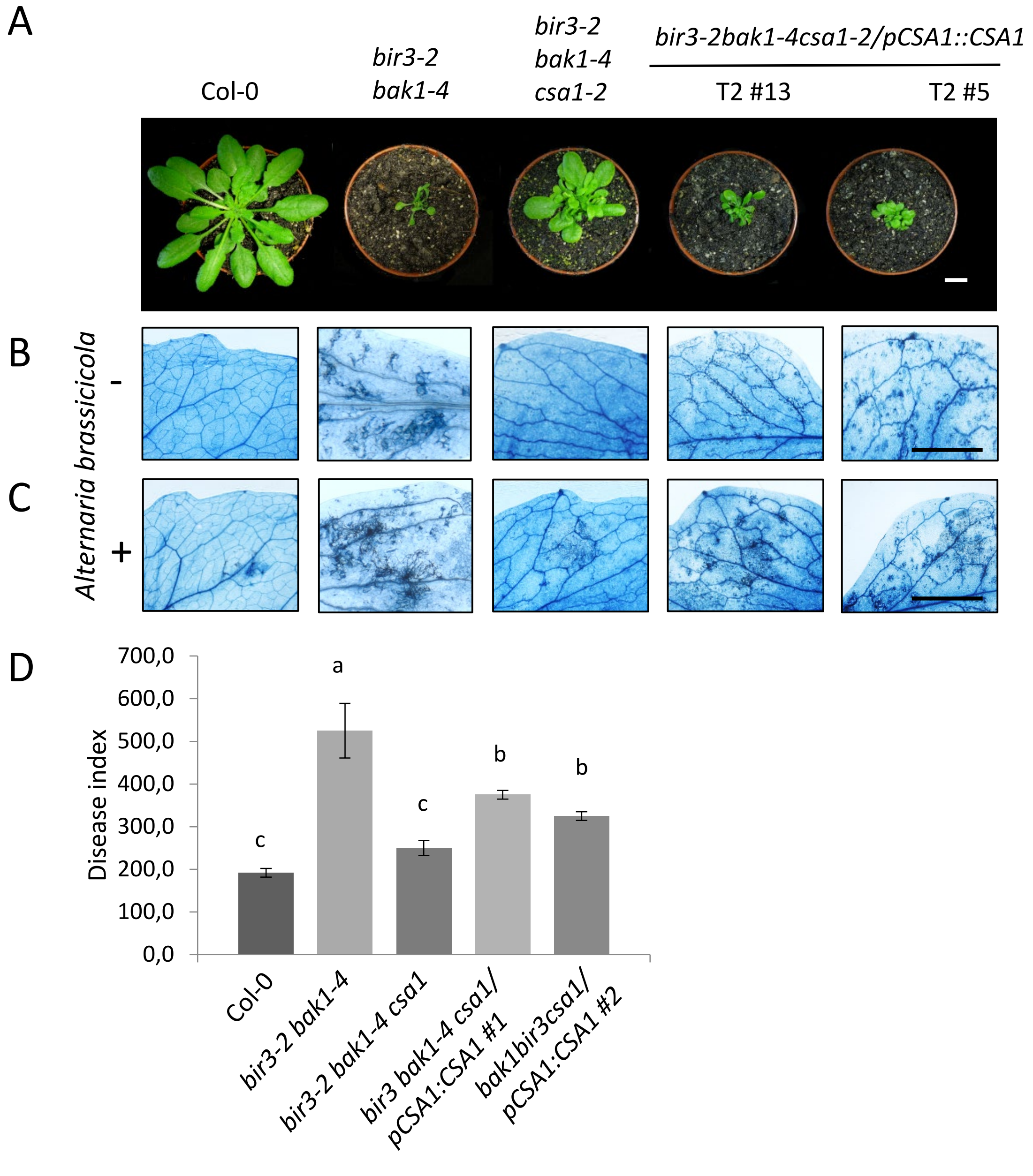


**Supplemental Figure 12: Expression of CSA1 can complement the *bak1 bir3 csa1* triple mutant phenotype**

**(A)** Representative pictures of the morphological phenotype of 6-week-old Col-0, *bir3-2 bak1-4,* double mutant, the triple mutant *bak1-4 bir3-2 csa1-2* and the complementation lines expressing *CSA1* under the endogenous promotor in the triple mutant background. The scale bar represents 1 cm. **(B)** uninfected leaves of the genotypes shown in (A) stained with trypan blue for cell death. **(C)** Leaves of the same genotypes as in (A) and (B) droplet-infected with *Alternaria brassicicola* and trypan blue stained. The scale bar in (B) and (C) represents 5 mm. **(D)** Disease indices of *Alternaria brassicicola* infected leaves of the indicated genotypes 13 days after infection shown as mean ± SE (n=12). Different letters indicate significant differences according to one-way ANOVA and Tukey's HSD test ($p<0.05$). The experiments were repeated at least three times with similar results.

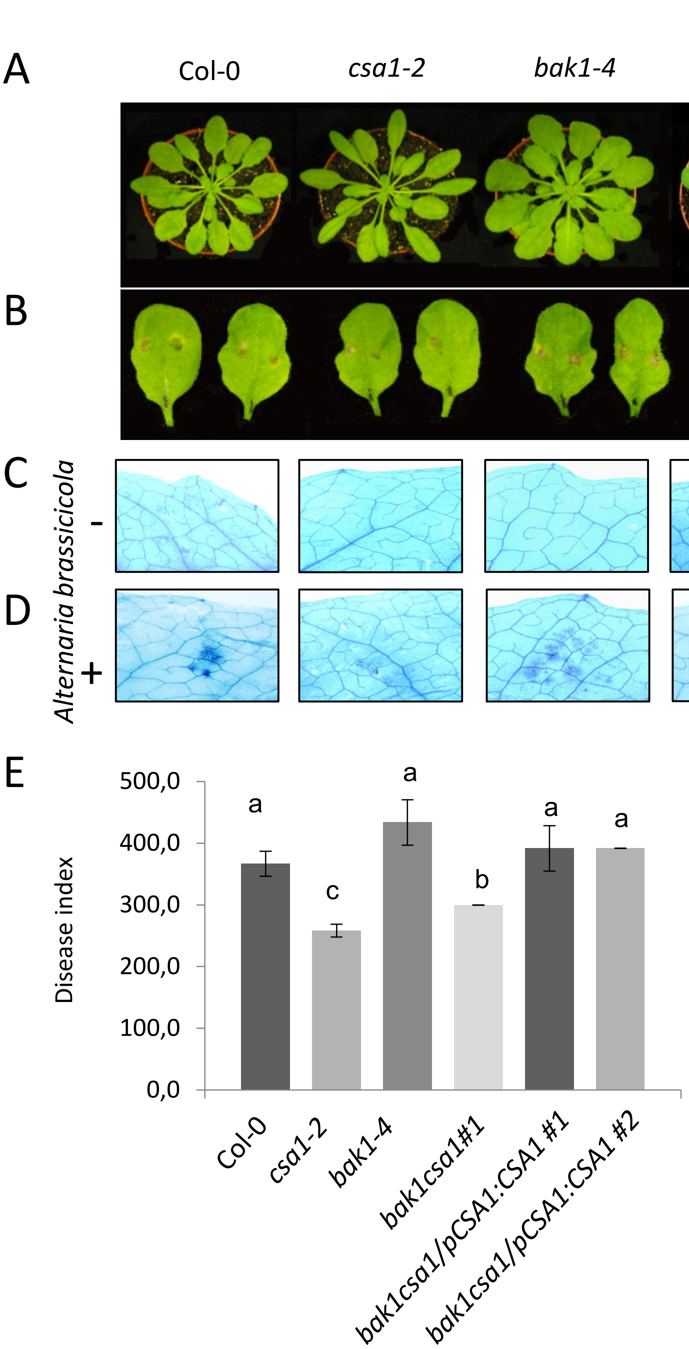


**Supplemental Figure 13: Expression of CSA1 can complement the *bak1 csa1* double mutant phenotype**

**(A)** Representative pictures of the morphological phenotype of 6-week-old Col-0, *csa1-2*, *bak1-4, bak1-4 csa1-2* and the complementation lines expressing CSA1 under the endogenous promotor in the double mutant background. **(B)** Leaves of the same genotypes as in A and B droplet-infected with *Alternaria brassicicola.* The scale bars in (A) and (B) represent 10 mm **(C)** Uninfected leaves of the genotypes shown in (A) stained with trypan blue for cell death. **(D)** Leaves of the same genotypes as in A droplet-infected with *Alternaria brassicicola* and trypan blue stained. The scale bars in (C) and (D) represent 5 mm. **(E)** Disease index of *Alternaria brassicicola* infected leaves of the indicated genotypes 13 days after infection shown as mean ± SE (n=12). Different letters indicate significant differences according to one-way ANOVA and Tukey's HSD test ($p<0.05$). The experiments were repeated at least three times with similar results.

**Supplemental Table 2: Primers used in this study**

| Name | Sequence (5’-> 3') | Characteristics |
|---|---|---|
| CSA1-F (KpnI) | CGGGTACCTAATGACAAGCTCCTCCTCCTG | cloning for CSA1 cds, fwd |
| CSA1-R (SalI) | AGGGTCGACATGGCTATACATTTCATAAAG | cloning for CSA1 cds, rev |
| BIR3_F (KpnI) | CGGGTACCATGAAGAAGATCTTCATCACAC | cloning for PCL-BIR3 or PNL-BIR3, fwd |
| BIR3_R (SalI) | AGGGTCGACAGCTTCTTGTTTGTTGAAGACC | cloning for PCL-BIR3 or PNL-BIR3, rev |
| CHS3-F (KpnI) | CGGGTACCATGGAACCACCAGCTGCTCG | cloning for CHS3 cds, fwd |
| CHS3-R (SalI) | AGGGTCGACTAACTTTGAATATTGTGGAG | cloning for CHS3 cds, rev |
| pXNubA22-F | CAAGCATACAATCAACTC | yeast cloning PCR, fwd |
| pXNubA22-R | ATTGATCCACCTCCACCG | yeast cloning PCR, rev |
| pMetYC-F | ATTCTATTACCCCCATCC | yeast cloning PCR, fwd |
| pMetYC-R | ATCCACCTCCACCGGATC | yeast cloning PCR, rev |
| CSA1-geno-F | CGGGTACCTAcacaattccagcatccacttgcg | cloning CSA1 genomic DNA, fwd |
| CSA1-geno-R | AGGGTCGACATGGCTATACATTTCATAAAGC | cloning CSA1 genomic DNA, rev |
| CHS3-geno-F | CGGGTACCATGGAACCACCAGCTGCTCG | cloning for CHS3 genomic DNA, fwd |
| CHS3-geno-R | ACCGCTCGAGTAACTTTGAATATTGTGGAGTCTTGG | cloning for CHS3 genomic DNA, rev |
| USER-CSA1-F | GGCTTAAUATGACAAGCTCCTCCTCCTGGGT | cloning CSA1, fwd |
| USER-CSA1-R | AACCCGAUCCACACAAAAGAGTGGAACCAAAACCAG | cloning CSA1, rev |
| USER-CHS3-F | GGCTTAAUATGGAACCACCAGCTGCTCGTG | cloning CHS3, fwd |
| USER-CHS3-R | AACCCGAUCCTAACTTTGAATATTGTGGAGTC | cloning CHS3, rev |
| USER-V5-F | ATCGGGTUCGCATTCGGGTAAGCCAATCCC | cloning of CSA1 and CHS3, fwd |
| USER-V5-R | GGTTTAAUAAGCTTAGGTTGAGTCGAGTCCGAG | cloning of CSA1 and CHS3, rev |

**Supplemental Table 2: Primers used in this study continued**

| | | |
|---|---|---|
| pad4-1-PflmI-F | ATGAGTCGCATAAGACTAGCCAAG | genotyping for *pad4-1* |
| pad4-1-PflmI-R | CCATTTCTTTCCTAAATGAAAATCA | genotyping for *pad4-1* |
| FP-sag101 | GATCTTGGAGATACATAACCC | genotyping for sag101-2 |
| BF53 | ACTTCCGGGTGTTCATAAACTCGGTCAAG | genotyping for sag101-2 |
| dSpm1 | CTTATTTCAGTAAGAGTGTGGGGTTTTGG | genotyping for sag101-2 |
| chs3-3 LP | ATTTTGAGCAGCTTCCTAGGC | genotyping *chs3-3*, fwd |
| chs3-3 RP | TCCTCATGATCTTTGGAATGC | genotyping *chs3-3*, rev |
| NDRF | GACGAGATTGCTCATTGCCATTGG | genotyping *ndr*1, fwd |
| NDRR | TAGGCATGGTACAATAACCGGAACC | genotyping *ndr1*, rev |
| js1259 | CTGGTTTCCACTTCACGATGA | genotyping for *eds1-12* |
| js959 | AACTAGCATACAGAGGGGCA | genotyping for *eds1-12* |
| js960 | GCTGAGAGAAATCGAACCGG | genotyping for *eds1-12* |
| At1g27190F3 | CTCGCCGGTGAGATTCCTGAGTCTCTTA | genotyping *bir3-2* , fwd |
| At1g27190R3 | ACAGACAAAGGCTTTTGCCCTGTAACCA | genotyping *bir3-2* , rev |
| bak1-4_F | CATGACATCATCATCATTCGC | genotyping for *bak1-4*, fwd |
| bak1-4_R | ATTTTGCAGTTTTGCCAACAC | genotyping for *bak1-4*, rev |
| csa1-2_LP | CATCCAGGAAAGCTAGTGCAG | genotyping for *csa1-2*, fwd |
| csa1-2_RP | GGCTGAAATTCCCGTTAAAAG | genotyping for *csa1-2*, rev |
| FRK1_fwd | TGCAGCGCAAGGACTAGAG | qRT-PCR of *FRK1* |
| FRK1_rev | ATCTTCGCTTGGAGCTTCTC | qRT-PCR for *FRK1* |
